# A single-domain antibody targets aggregation-prone region of *α*-synuclein to reduce synucleinopathy, rescue neurodegeneration and improve function

**DOI:** 10.64898/2026.07.07.735601

**Authors:** Pragati, Erin E Congdon, Yixiang Jiang, Hediye Erdjument-Bromage, Huai-Wei Huang, Ruimin Pan, Isabella S Marchal, Xiang-Peng Kong, Thomas A. Neubert, Hyung Don Ryoo, Einar M Sigurdsson

## Abstract

Synucleinopathies are a group of neurodegenerative disorders characterized by the accumulation of aggregated α-synuclein (α-syn), including Parkinson’s disease, Dementia with Lewy Bodies, and Multiple System Atrophy. These diseases are marked by locomotor and non-motor impairments, as well as mitochondrial dysfunction and the loss of dopaminergic (DA) neurons. We have developed several anti-α-syn single-domain antibodies (sdAbs) and demonstrated the diagnostic imaging potential of two of them and the acute therapeutic benefit of one in clearing α-syn in a mouse model. However, whether these sdAbs can suppress α-syn-mediated neuronal loss and locomotor impairment in vivo remains unclear. We evaluated the therapeutic potential of five anti-α-syn sdAbs to clear pathological α-syn in mouse neuronal culture and then demonstrated their in vivo efficacy in a *Drosophila* model of synucleinopathy. The sdAbs differed in their efficacy to lower levels of phospho-serine 129 α-syn, prevent loss of DA neurons, alleviate mitochondrial dysfunction, improve motor function, and prolong survival in synucleinopathy flies. The most effective sdAb, 2H1, has not been reported before. It binds strongly to the aggregation prone region of α-syn and robustly improves all these disease parameters. Additionally, that sdAb is associated with α-syn in the fly neurons, as shown through proximity dependent turboID biotinylation assays. The sdAb-turboID also biotinylated α-syn-associated proteins involved in synapse/vesicle trafficking pathways, pinpointing the location of their intracellular interaction. Our findings provide an insight into the therapeutic mechanism of action of these sdAbs and strongly support their clinical development.

## Introduction

Aggregates of α-synuclein (α-syn), such as Lewy bodies (LBs), Lewy neurites, and glial cell inclusions, are linked to a group of neurodegenerative diseases known as synucleinopathies. These include Parkinson’s disease (PD), Dementia with Lewy bodies (DLB), and Multiple System Atrophy [MSA; (*1*)]. These diseases are characterized by motor symptoms, such as tremor, rigidity, and bradykinesia, as well as non-motor symptoms, including sleep disturbances, hyposmia, and certain neuropsychiatric issues (*2*).

α-syn is a small (14 kDa) acidic protein found in neurons of both the central and peripheral nervous systems and other tissues (*3*). Endogenous α-syn mainly exists as a folded tetramer (∼58 kDa) with little tendency to form amyloid aggregates. It has three main regions with distinct functions: amino acids 1–60 form the N-terminus with amphipathic repeats for membrane interactions, residues 61–95 constitute the hydrophobic aggregation-prone region, and the C-terminus (96–140) is negatively charged, involved in Ca^2+^ binding and chaperone-like activities (*4*). Post-translational modifications, such as phosphorylation of Ser129 in the C-terminal region, affect α-syn folding and aggregation (*5*). Normally, less than 4% of soluble α-syn is phosphorylated. However, about 90% of α-syn in LBs is phosphorylated, highlighting a link between Ser129 phosphorylation and α-syn aggregation in synucleinopathies (*6–8*). Additionally, aggregated α-syn in LBs has been linked to mitochondrial dysfunction (*9, 10*). Specifically, oligomeric/phosphorylated α-syn alters mitochondrial respiration (*11*) and oxidizes ATP synthase, causing mitochondrial swelling and cell death (*12*). It is also a critical determinant of mitochondrial dysfunction in dopaminergic neurons (*13*).

Familial forms of synucleinopathies are linked to mutations or duplications/triplications of the SNCA gene that codes for α-syn, demonstrating its role in disease pathogenesis and a key therapeutic target. Immunotherapies targeting α-syn have shown promise in preclinical models. There are several α-syn immunotherapies in various phases of clinical trials (https://www.alzforum.org/therapeutics). These clinical IgG antibodies face challenges in crossing the blood-brain barrier (BBB) due to their large size (∼150 kDa). Smaller antibody fragments, such as single domain antibodies [sdAbs, variable heavy domain of heavy chain only antibodies (VHH)] occur naturally in camelids and are much smaller than traditional IgG antibodies (∼15 kDa vs. ∼150 kDa) (*14*). Their small size enhances brain penetration, enables binding to hidden epitopes, and facilitates intracellular folding, making them suitable for gene therapy. A key disadvantage compared to IgGs is their short half-life, though this is less challenging in gene therapy and is helpful for diagnostic imaging. We recently developed several sdAbs against human α-syn by immunizing a llama with full-length recombinant α-syn and have reported on the suitability of two of them as diagnostic imaging agents in a mouse model of synucleinopathy (*15*). Further, we have shown that one of the sdAbs, fused with a protein degrader, promotes proteasomal degradation of α-syn in primary cultures and in a mouse model of synucleinopathy (*16*). However, whether these sdAbs can prevent α-syn-mediated neuronal loss and functional impairments in vivo remains unclear.

In view of the limited in vivo evidence demonstrating the therapeutic potential of these anti-α-syn sdAbs, we evaluated five anti-α-syn sdAbs by transgenically expressing them in synucleinopathy flies. Our findings show that these sdAbs exhibit varying efficacy in mitigating motor deficits, reducing pathogenic α-syn phosphorylation, prolonging lifespan, and limiting loss of dopaminergic (DA) neurons. The most efficacious sdAb, 2H1, rescued all these characteristics associated with synucleinopathy. Additionally, 2H1 improved mitochondrial function and interacted with α-syn and its associated proteins in vivo, particularly with proteins involved in synapse and vesicle-trafficking pathways, in α-syn-expressing flies as indicated by proximity-dependent biotinylation-mediated interactions. Taken together, we propose that anti-α-syn sdAbs confer therapeutic benefits by counteracting various toxic effects mediated by aggregated α-syn.

## Results

### Anti-*α*-syn sdAbs clear *α*-syn and prevent extra- and intracellular toxicity of *α*-syn in a neuronal culture model

To determine the efficacy of anti-α-syn sdAbs, we first tested them in a primary neuronal culture model prepared from transgenic (tg) M83 mice. These animals express the familial A53T α-syn mutation, to which a toxic α-syn fraction (S1) from a human synucleinopathy patient was added to enhance pathology and to better relate the model to the human condition. As described in the Method section, to simulate extracellular (α-syn + sdAb) and intracellular (α-syn → sdAb) targeting of pathological α-syn, neurons cultured from postnatal day 0 were treated with 10 µg/ml of S1 α-syn brain supernatant, with or without sdAb (1 µg/ml for 7 days).

In the intracellular paradigm (Fig. 1A-B), α-syn alone was toxic compared to untreated control (46% decrease in GAPDH, p <0.0001), and this toxicity was prevented by sdAbs 2D10 (86% of control; p = 0.0160), 2H1 (106% of control, p <0.0001), and 1G10 (172% of control, p <0.0001; Fig 1A). The human brain α-syn fraction led to a 5.5-fold increase in α-syn levels in the neurons (p = 0.0003; Fig. 1B), which was prevented in all sdAb-treated cells except for 1D12 (α-syn/GAPDH ratios: 0.50, 1.00, 1.43, 1.34, and 0.63 compared to untreated control set at 1; p = 0.0009, 0.0034, 0.0103, 0.0081, and 0.0013 for sdAbs 2D8, 2D10, 2H1, 2H7, and 1G10, respectively; Fig. 1B). Representative and complete blots used for quantifications are shown in Fig. S1A and S2.

**Fig. 1.**
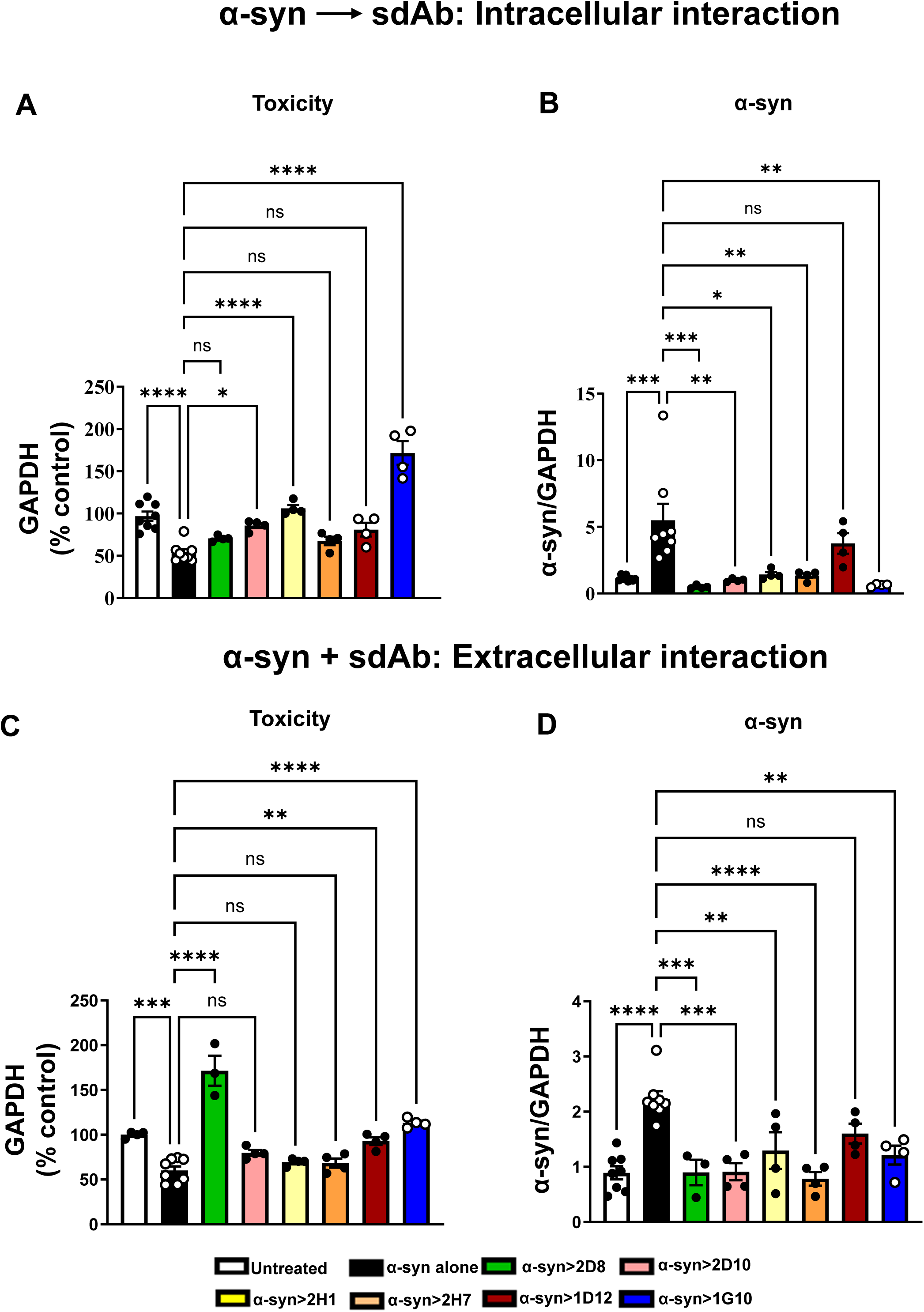
Anti-*α*-syn sdAbs clear *α*-syn and prevent toxicity of *α*-syn-enriched fraction from human synucleinopathy brain in primary tg A53T *α*-syn mouse neurons. (A-D) Primary neuronal cultures from M83 A53T α-syn mice collected at postnatal day 0 were exposed to 10 µg/ml of S1 α-syn brain supernatant with or without sdAb (1 µg/ml for 7 days). Each sdAb was administered: 1 day later after media wash to remove remaining extracellular α-syn that was not taken up into the neurons (intracellular interaction; A-B) or at the same time as the S1 α-syn fraction (extracellular interaction; C-D). α-syn alone was toxic compared to untreated control (white compared to black, 46% reduction). Three of the sdAbs, 2D10 (pink), 2H1 (yellow), and 1G10 (blue), prevented its intracellular toxicity (86, 106, and 172% of control, respectively) as demonstrated by increased GAPDH levels (A). The α-syn fraction seeded α-syn aggregation within the neurons (5.5-fold above untreated control neurons), which was prevented in all the sdAb treated cells except 1D12 (brown) (B). α-syn alone was toxic compared to untreated control (40% reduction). Three of the sdAbs, 2D8 (green), 1D12 (brown), and 1G10 (blue), prevented this toxicity extracellularly (171%, 93%, and 113% of control) as demonstrated by increased GAPDH levels (C). As in B, α-syn seeded aggregation within the neurons (2.2-fold above untreated control cells), which was prevented in all the sdAbs treated neurons except 1D12 (brown) (D). Bar graphs are presented as mean ± SEM from 4–8 replicates. One-way ANOVA, Tukey’s multiple comparisons test. *p ≤ 0.05; **p ≤ 0.01, *** p ≤ 0.001 ****p ≤ 0.0001 and ns = non-significant.

In the extracellular paradigm (Fig. 1C-D), α-syn alone was also toxic and decreased GAPDH levels by 40% (p = 0.0002), and this toxicity was prevented by sdAbs 2D8, 1D12, and 1G10 (171%, 93% and 113% of untreated control, p <0.0001, p = 0.0021, and p <0.0001; Fig. 1C). Here, the human brain α-syn fraction also increased α-syn levels (2.2-fold above control, p <0.0001, Fig. 1D), which was prevented by all anti-α-syn sdAbs except 1D12 (α-syn/GAPDH ratios: 0.89, 0.91, 1.3, 0.78, and 1.2, compared to untreated control set at 1; p = 0.0004, 0.0001, 0.0093, <0.0001 and 0.0038 for sdAbs 2D8, 2D10, 2H1, 2H7 and 1G10, respectively; Fig. 1D). Representative and complete blots used for quantifications are shown in Fig. S1B and S3. These results demonstrate the varying in vitro efficacy of different α-syn sdAbs in clearing α-syn and preventing toxicity in mouse neurons.

### Anti-*α*-syn sdAbs suppress climbing defects associated with synucleinopathy in female flies

To assess the in vivo therapeutic effectiveness of anti-α-syn sdAbs, we created five transgenic fly lines expressing these sdAbs except 1G10: UAS-2D8, UAS-2D10, UAS-2H1, UAS-2H7, and UAS-1D12. Additionally, for thorough analysis, we developed a transgenic fly expressing an sdAb that does not bind to α-syn but targets an unrelated epitope of the dengue virus (UAS-Dv^sdAb^), serving as a negative or extra UAS control (*17, 18*). All these transgenes include a histidine tag and a myc tag and were inserted at the same genomic locus. These sdAbs did not contain secretory signal peptides, designed to be expressed only in the cytoplasm. We expressed these sdAbs in all neurons using *Elav-Gal4* and assessed their levels through western blot utilizing different antibodies (Fig. S4A-B; for the full blot see Fig. S6). First, we determined the expression levels of sdAbs with an anti-myc tag antibody. 1D12 was not detected and 2D8 gave a faint band, 2H7 had a moderate band whereas 2D10 and 2H1 had the strongest bands (Fig. S4A). We also evaluated the expression of these sdAbs using an anti-VHH antibody. Here, 1D12 revealed a faint band, 2D8 and 2H7 had moderate bands, whereas 2D10 and 2H1 had the strongest bands (Fig. S4B). Hence, all the sdAbs are expressed in the flies, but their expression levels or accessibility of their detection epitopes may differ. For example, the myc epitope is not always accessible in proteins (*19*). Complete blots for Fig. S4 are provided in Fig. S6. Additionally, we analyzed the transcript levels of these sdAbs and observed significant levels of sdAbs 2D8 (p = 0.0002), 2D10 (p = 0.0002), and 2H1 (p = 0.0013) relative to 2H7. Interestingly, sdAb 1D12 had the highest transcript level (p <0.0001) but the lowest apparent protein expression (Fig. S4C), suggesting low stability, differences in translation efficiency as the transgenes were inserted at the same genetic locus, or perhaps differences in exposure of their detection epitopes.

After confirming their expression (transcription and translation), we performed a climbing assay to assess the therapeutic benefits of different sdAbs targeting α-syn. We specifically measured negative geotaxis, where we assessed the upward movement of flies that were tapped to the bottom of the vial (see Materials and Methods). Expression of α-syn in flies causes locomotor deficits, reducing their climbing ability (*20, 21*). We specifically found that pan-neuronal expression of human α-syn (*Elav>α-syn*) decreased the climbing ability of control flies from 86% to 43% in 10-day-old adult male flies (Fig. S5A). This decline further progressed to 27%, 6%, and 2% in 15, 20, and 25-day-old males, respectively (Fig. S5B-D). *Elav>α-syn+Dv^sdAb^* flies showed a similar progressive decline in climbing ability as *Elav>α-syn* flies (Fig. S5A-D). Pan-neuronal co-expression of anti-α-syn sdAbs did not rescue the climbing defects in adult male flies.

We also analyzed the effect of anti-α-syn sdAbs in adult female flies. As noted in the males, pan-neuronal expression of α-syn in females also decreased the climbing ability of control flies, specifically from 80% to 44% in 10-day-old adult female flies (Fig. 2A), which further declined to 37%, 13%, and 9% in 15, 20, and 25-day-old females, respectively. A comparable pattern of decline was seen in the control α-syn+Dv^sdAb^ flies. Pan-neuronal co-expression of anti-α-syn sdAbs did not improve the climbing ability of 10-day-old female flies (Fig. 2A), but 15-day-old females showed significant improvement (2D8: p = 0.0256; 2D10: p = 0.0007; 2H1: p<0.0001; 2H7: p = 0.0002; and 1D12: p = 0.0146; Fig. 2B).

**Fig. 2.**
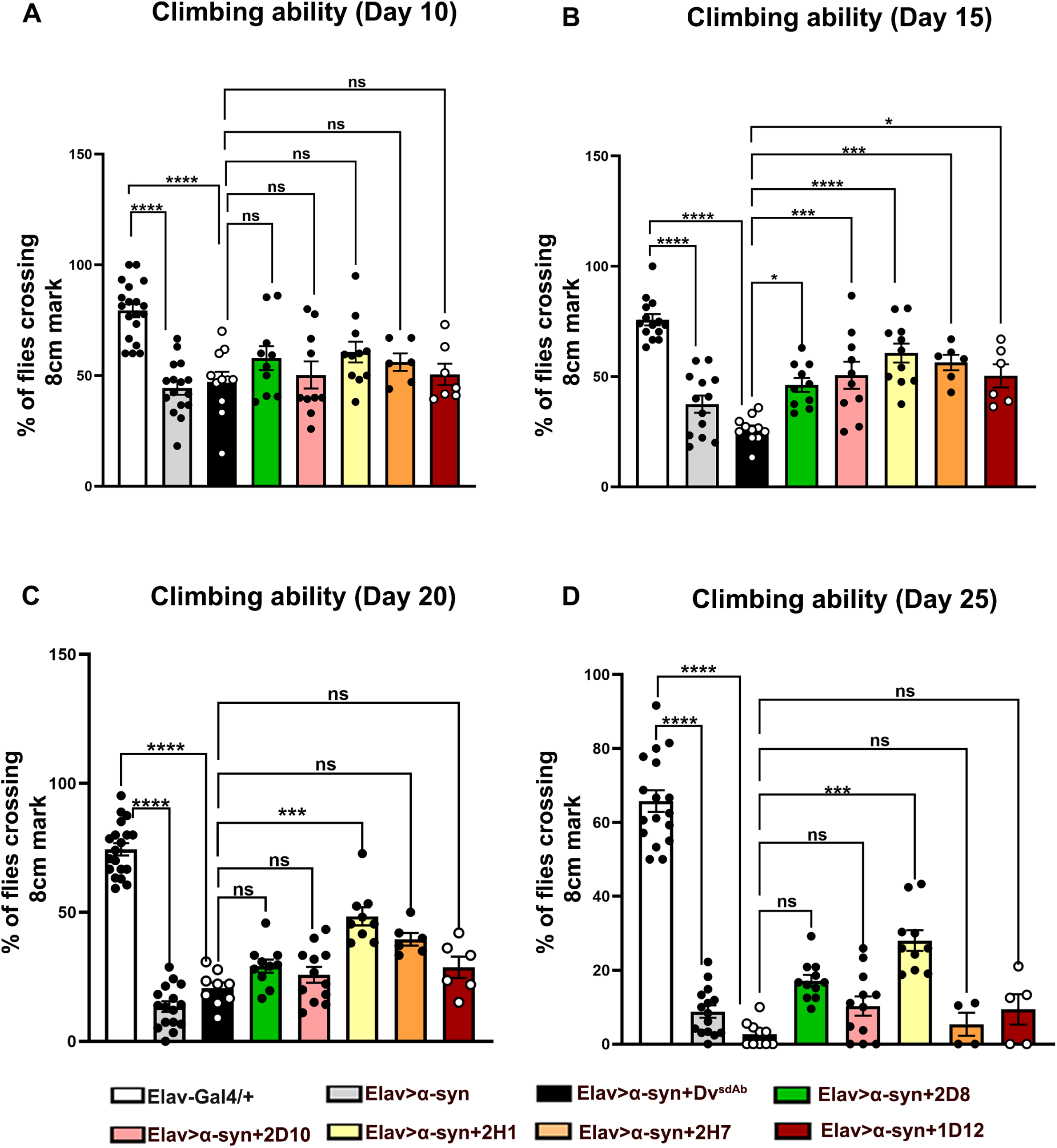
Pan-neuronal expression of anti-*α*-syn sdAbs reduces climbing defects in *α*-syn-expressing female flies. (A-D) Bar graphs showing a quantitative assessment of the relative climbing ability of different age-matched adult female flies. Results from day 10 (A), day 15 (B), day 20 (C), and day 25 (D) are shown. Different genotypes are marked with distinct colors as indicated in the figure legend. Flies not expressing α-syn (white bars) climbed the 8 cm bars more quickly than those expressing α-syn. Grey and black bars show flies expressing α-syn and α-syn with a control sdAb (Dv^sdAb^). Those two groups had impaired motor function at all time points, which worsened with age. All five anti-α-syn sdAbs improved climbing ability at day 15 post-eclosion (B), compared to the control Dv^sdAb^ group, but only sdAb 2H1 remained effective at days 20 and 25 (C-D). Bar graphs are presented as mean ± SEM, and each data point represents an average of 10-22 flies in a vial. Genotypes and the number of flies analyzed per group: *Elav-Gal4/+* (N = 175), *Elav>α-syn* (N = 194), *Elav>α-syn+Dv^sdAb^*(N = 112), *Elav>α-syn+2D8* (N = 121), *Elav>α-syn+2D10* (N = 100), *Elav>α-syn+2H1* (N = 120), *Elav>α-syn+2H7* (N = 100), and *Elav>α-syn+1D12* (N = 100). Two-way ANOVA, Tukey’s multiple-comparison test. *p ≤ 0.05, ***p ≤ 0.001, ****p ≤ 0.0001 and ns = non-significant.

The sdAb-mediated functional improvement in females and not in males may be because our Elav-Gal4 is located on the X chromosome. This effectiveness persisted for sdAb 2H1 at 25 days post-eclosion (day 20: p = 0.0002; day 25: p = 0.0006; Fig. 2C and D). These results indicate varying efficacy of different anti-α-syn sdAbs in mitigating α-syn-induced locomotor impairments, with 2H1 being the most effective. We note a correlation between 2H1’s effectiveness and its high protein expression level.

### Anti-*α*-syn sdAbs prevent *α*-syn-mediated degeneration of DA neurons in flies

Given the vulnerability of dopaminergic (DA) neurons to degeneration in synucleinopathies (*22*), we next assessed the efficacy of the anti-α-syn sdAbs in preventing DA neuron loss as detected by tyrosine hydroxylase (TH) staining. This enzyme is involved in DA production (*23*). It is produced in the cell body and transported to the axon and presynaptic terminals, making it an effective marker for visualizing DA neurons (*24*). The adult *Drosophila* brain contains about 130–200 DA neurons arranged into roughly eight bilateral clusters, including the anterior groups PAM (protocerebral anterior medial), PAL (protocerebral anterior lateral), T1 (first thoracic segment/subesophageal), and Sb (subesophageal). Additionally, there are the posterior groups PPL1 (protocerebral posterior lateral 1), PPL2ab, PPL2c, and PPM1/2/3 (protocerebral posterior medial) (*25*). Compared to normal non-transgenic controls, which have all these DA neuronal clusters in 15-day-old adult female flies (Fig. 3A; see also I, J), pan-neuronal expression of α-syn significantly promoted degeneration of the PPL1 and PPM2 DA neuronal clusters (compare Fig. 3B with A; see also I, J). Our results align with previous studies reporting DA neuron loss in adult flies of this synucleinopathy model (*26, 27*). Like *Elav>α-syn* flies, pan-neuronal co-expression of *Dv^sdAb^* with α-syn also led to notable deterioration of the PPL1 and PPM2 clusters (compare Fig. 3C with B and A; see also I, J). In contrast, the loss of PPL1 was prevented by pan-neuronal co-expression of sdAbs 2D8 (p = 0.0390), 2D10 (p = 0.0301), and 2H1 (p = 0.0434), whereas sdAbs 2H7 and 1D12 were ineffective (compare PPL1 region in Fig. 3D-H with C; see also I, J). Likewise, degeneration of the PPM2 cluster was prevented by sdAbs 2D10 (p = 0.0225), 2H1 (p = 0.0012), and 1D12 (p = 0.0225), while sdAbs 2D8 and 2H7 were ineffective (compare PPM2 region in Fig. 3D-H with C; see also I, J). In addition, we assessed the impact of these sdAbs in *Elav>α-syn* flies as they age and noted that sdAb 2H1 (p = 0.0220) rescued the loss of the PPL1 neuronal population in 25-day-old female flies (compare PPL1 region in Fig. S7F with C; see also I, J). Similarly, degeneration of the PPM2 cluster was prevented by sdAbs 2D8 (p = 0.0389; compare PPM2 region in Fig. S7D with C; see also I, J). These results indicate that anti-α-syn sdAbs differ in their ability to prevent degeneration of DA neurons in a fly model of synucleinopathy. Interestingly, sdAbs 2D8, 2D10, 2H1, and 1D12 consistently improved not only motor function but also neuronal survival, with 2H1 and 2D8 remaining effective in aged animals.

**Fig. 3.**
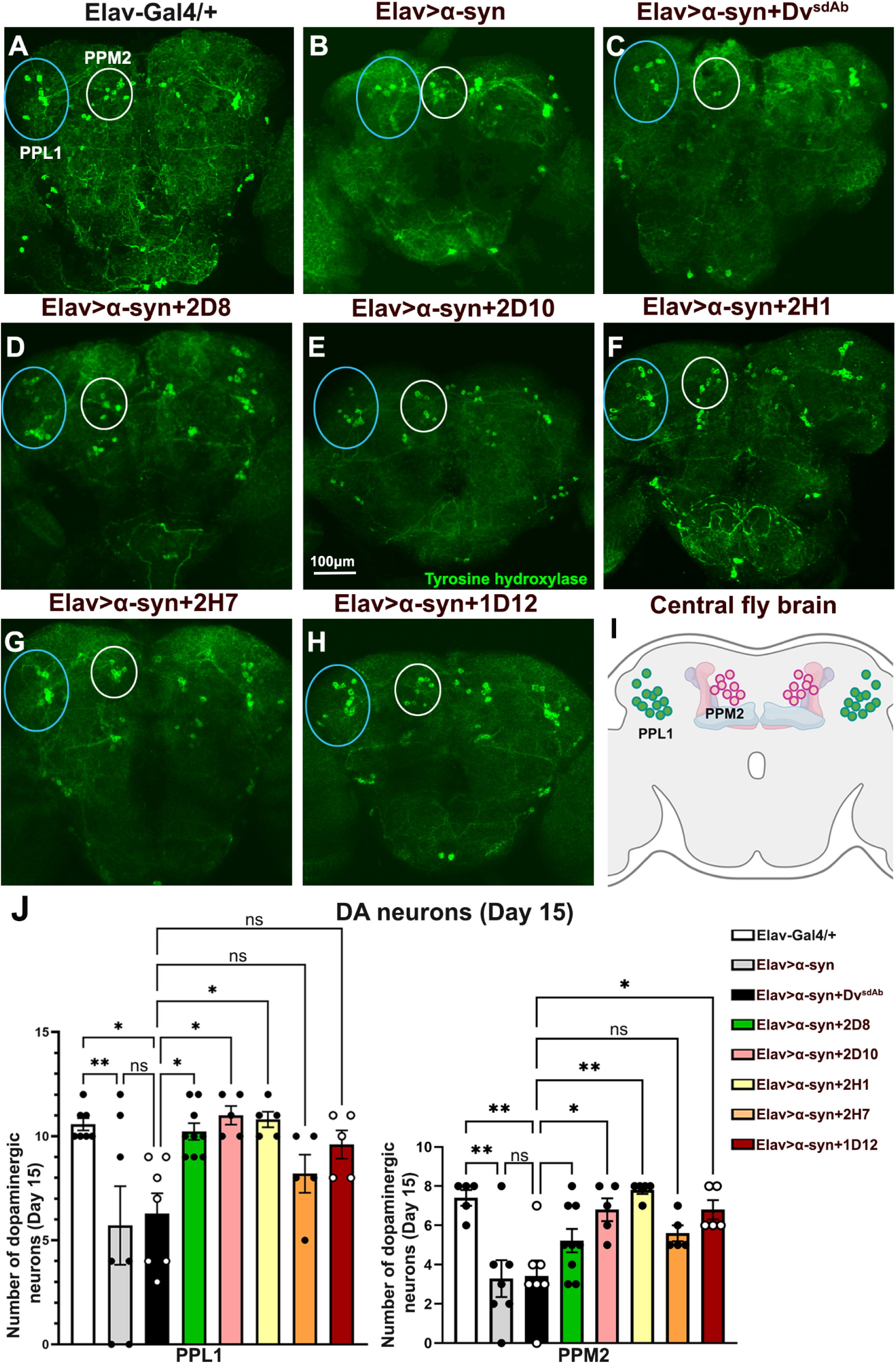
Neuron-specific expression of anti-*α*-syn sdAbs prevents the loss of DA neurons in *α*-syn-expressing flies. (A-H) Confocal images of a 15-day-old female adult brain stained with anti-tyrosine hydroxylase antibody. The marked circles represent PPL1 (blue) and PPM2 (white) neuronal clusters. (I) The schematic shows PPL1 and PPM2 neuronal clusters in an adult fly brain. (J) The average number of DA neurons in PPL1 and PPM2 neuronal clusters in different age-matched female adult brains per hemisphere (N = 5-9 per genotype). Quantification shows loss of DA neurons in *Elav>α-syn* and *Elav>α-syn+Dv^sdAb^* (control sdAb), which was prevented by all sdAbs except 2H7 and 1D12 in PPL1 and 2D8 and 2H7 in PPM2 neuronal clusters. Bar graphs are presented as mean ± SEM. One-way ANOVA, Tukey’s multiple-comparison test. *p ≤ 0.05, **p ≤ 0.01 and ns = non-significant; Scale bar A-H = 100 µm.

### Anti-*α*-synuclein sdAbs reduce pathological phosphorylated *α*-syn in flies

α-syn undergoes several post-translational modifications in synucleinopathies, including ubiquitination, phosphorylation, and nitration (*28*). Among these, phosphorylation of α-syn is significantly associated with its aggregation and disease progression (*29*). Under normal conditions, less than 4% of α-syn is phosphorylated. In contrast, approximately 90% of α-syn in Lewy bodies is phosphorylated and serves as a useful biomarker, with p-Ser129 being the most prominent marker (*6–8, 30*). To determine whether the enhanced motor ability and decreased loss of DA neurons observed in synucleinopathy flies expressing anti-α-syn sdAbs are related to sdAb-mediated reduction of pathological α-syn, we measured the levels of α-syn p-Ser129 and total α-syn in 15-day-old females. Western blot analysis showed that, compared to control flies that do not express α-syn and were immunonegative for α-syn p-Ser129 and α-syn, lysates from *Elav>α-syn* and *Elav>α-syn+Dv^sdAb^* showed significant presence of phosphorylated and total α-syn (Fig. 4A, rows 1-2, lanes 1-3; see also B and C). Compared to *Elav>α-syn+Dv^sdAb^*, sdAbs 2D8 and 2D10 did not alter these levels (compare Fig. 4A, row 1, lanes 4-5 to lane 3; see also B), while sdAbs 2H1 (p = 0.0031), 2H7 (p = 0.0044), and 1D12 (p = 0.0116) reduced α-syn p-Ser129 levels (compare Fig. 4A, row 1, lanes 6-8 to lane 3; see also B). However, expression of anti-α-syn sdAb in synucleinopathy flies did not change total α-syn levels (Fig. 4A, row 2; see also C). This highlights the specificity of these sdAbs in reducing pathological α-syn levels. Additionally, the neuronal marker Elav did not differ between groups, indicating the absence of global neuronal loss in any group (Fig. 4A, row 3; see also D). Complete blots for Fig. 4 are provided in Fig. S8.

**Fig. 4.**
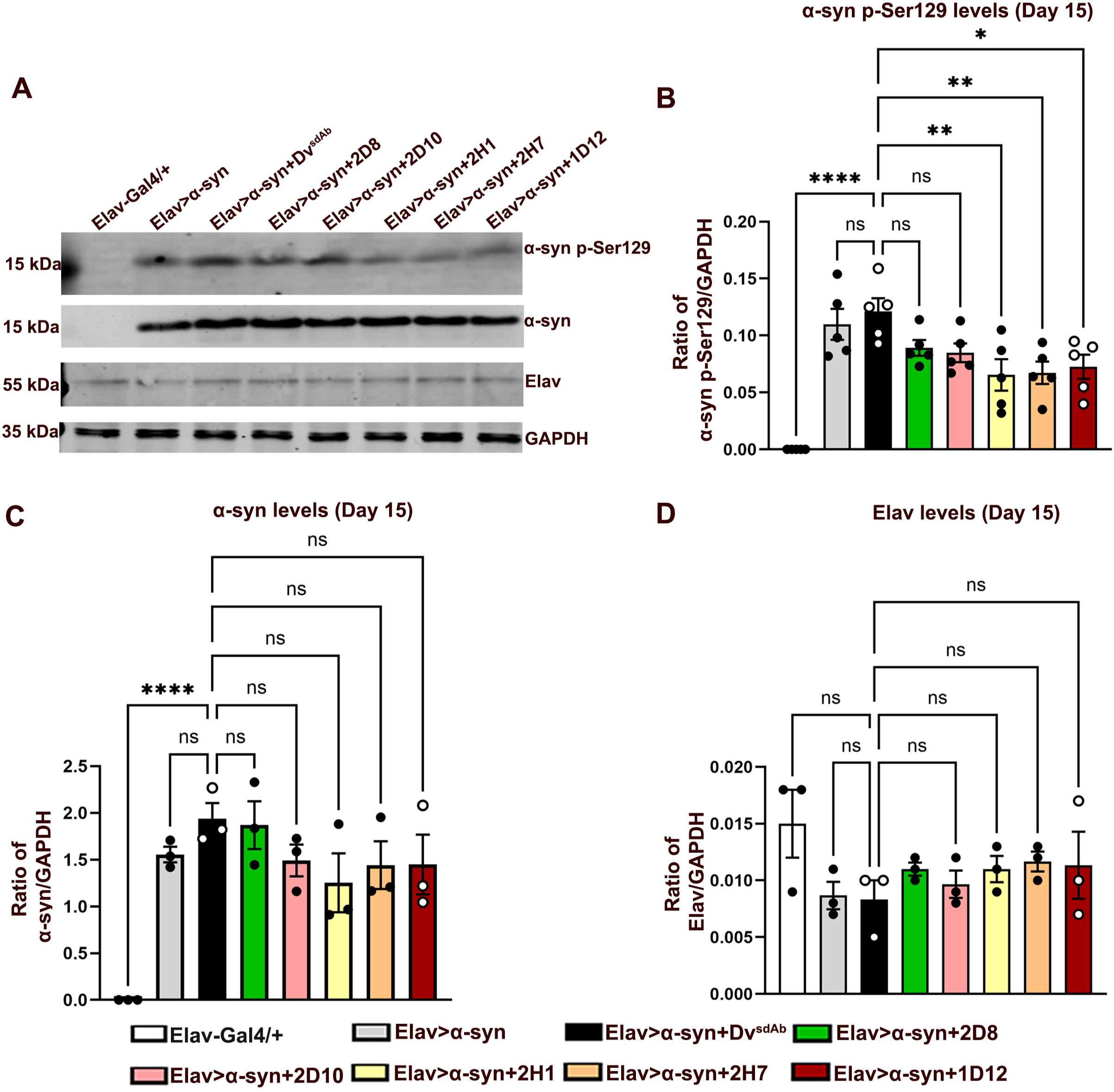
Neuronal expression of anti-*α*-syn sdAbs reduces levels of pathological *α*-syn p-Ser129 in *α*-syn flies. (A) Representative immunoblots of total cell lysate from 15-day-old adult female brains (N = 30 per biological replicate) show levels of α-syn p-Ser129, total α-syn, pan neuronal marker (Elav), and loading control GAPDH across different genotypes. (B-D) Quantification of α-syn p-Ser129 (B), total α-syn (C), and pan neuronal marker Elav (D) normalized to GAPDH levels in at least three independent biological replicates. sdAbs 2H1, 2H7, and 1D12 reduced α-syn p-Ser129 levels, but neither these nor other sdAbs altered total α-syn levels or the neuronal marker Elav. Bar graphs are presented as mean ± SEM. One-way ANOVA, Dunnett’s multiple comparisons test. *p ≤ 0.05, **p ≤ 0.01, ****p ≤ 0.0001 and ns = non-significant.

The western blot results were validated and extended by staining adult female brains with anti-α-syn p-Ser129. Compared to control brains, which showed no immunoreactivity, the adult brains of *Elav>α-syn* and *Elav>α-syn+Dv^sdAb^* flies displayed significant levels of phosphorylated α-syn, especially in the mushroom body and antennal lobe (see Fig. S9A-C; also refer to I, J). In comparison to the adult brains of *Elav>α-syn+Dv^sdAb^*, expression of sdAbs 2D8 and 2D10 (compare Fig. S9D and E with C; see also I, J) did not affect α-syn p-Ser129 levels, while expression of sdAbs 2H1 (p = 0.0366), 2H7 (p = 0.0035), and 1D12 (p = 0.0166; compare Fig. S9F-H with C; see also I, J) reduced α-syn p-Ser129 levels. These findings support the western blot data, and confirm that sdAbs 2H1, 2H7, and 1D12 effectively reduce pathological levels of phosphorylated α-syn.

### Anti-*α*-syn sdAbs also mitigate *α*-syn-mediated pathology in DA neurons

As shown above, pan-neuronal expression of α-syn causes degeneration of DA neurons, highlighting their vulnerability. Since co-expression of sdAbs reduced DA neuron loss, we expressed human α-syn only in DA neurons to directly assess the effectiveness of the sdAbs in these vulnerable neurons. We examined the levels of pathological α-syn p-Ser129 in DA neurons across different genotypes. Expression of α-syn solely in DA neurons led to robust accumulation of α-syn p-Ser129 in various DA neuron clusters (compare Fig. 5A with B-C; see also I, J). Compared to *TH>α-syn+Dv^sdAb^*adult brains, expression of sdAb 2D8 and 2D10 had no effect (compare Fig. D-E with C; see also I), whereas sdAbs 2H1 (p < 0.0001), 2H7 (p < 0.0001), and 1D12 (p < 0.0001) reduced the levels of pathological α-syn p-Ser129 within DA neurons (compare Fig. 5F-H with C; see also I). Additionally, co-staining of the brain with tyrosine hydroxylase antibody confirmed that the α-syn p-Ser129 accumulated in DA neurons across different genotypes (Fig. 5B’- H’). We also quantified the number of DA neurons in these genotypes. Compared to non-transgenic controls, expression of α-syn or co-expression of *Dv^sdAb^* with α-syn solely in DA neurons also led to notable deterioration of the PPL1 and PPM2 clusters (compare Fig. S10A with B-C; see also I, J). Interestingly, in comparison to the *TH>α-syn+Dv^sdAb^* adult brains, the loss of PPL1 clusters was prevented by pan-neuronal co-expression of sdAbs 2H1 (p = 0.0430) and 1D12 (p = 0.0052); however, sdAbs 2D8, 2D10, and 2H7 were ineffective (compare PPL1 region in Fig. S10D-H with C; see also J).

**Fig. 5.**
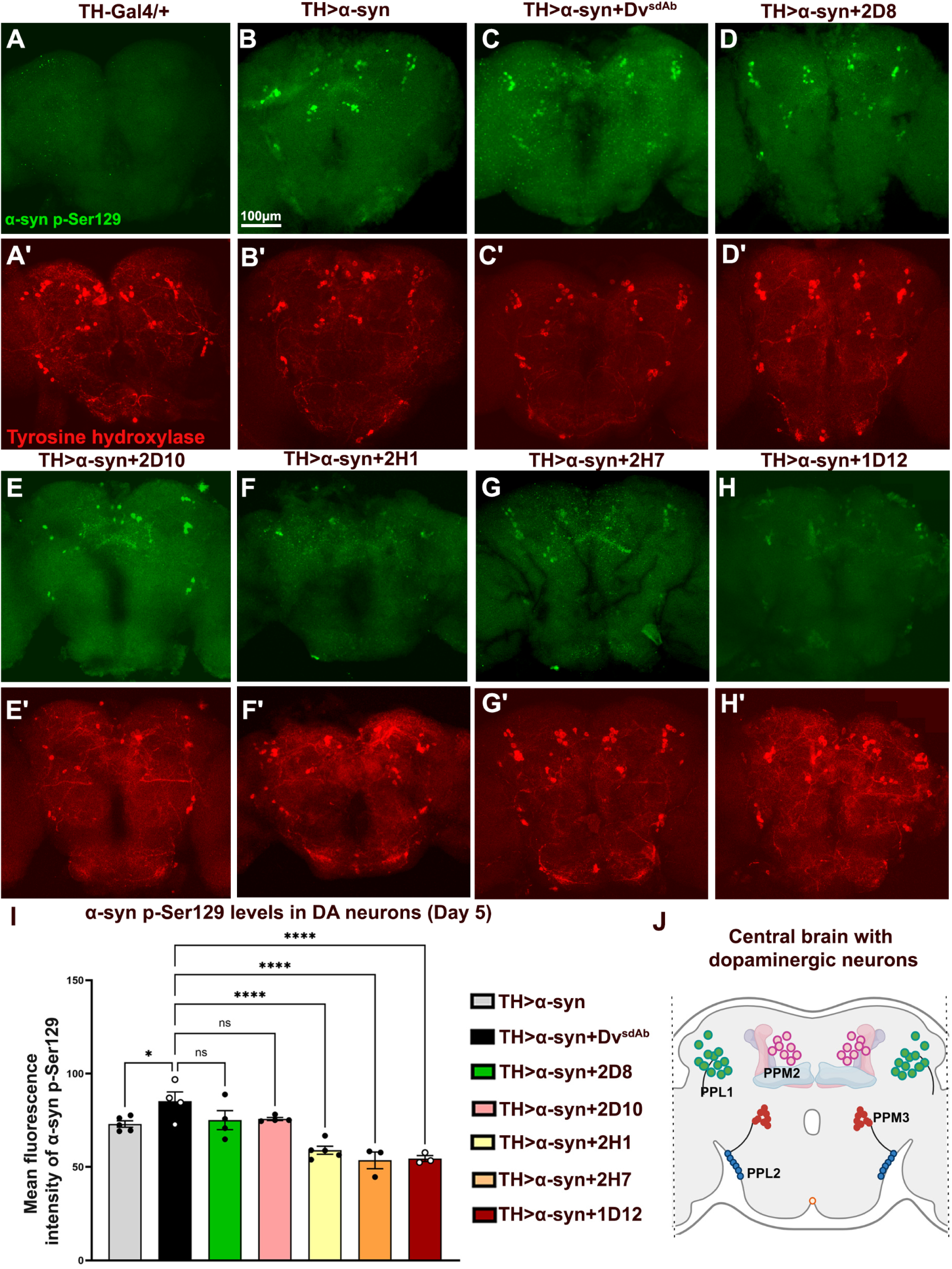
Exclusive expression of anti-*α*-syn sdAbs and *α*-syn in DA neurons reduces pathological *α*-syn p-Ser129 immunoreactivity in the brains of adult female flies. (A-H’) Confocal images of a 5-day-old female adult brain stained with anti-tyrosine hydroxylase (TH, a marker of DA neurons) and an anti-α-syn p-Ser129 antibody. *TH-Gal4/+* control flies were negative for α-syn p-Ser129 (A). Immunostained images of different genotypes stained with tyrosine hydroxylase confirmed detection of α-syn p-Ser129 immunoreactivity in DA neurons (B-H’). (I) Quantification of the fluorescence intensity of α-syn p-Ser129 in DA neurons shows that sdAbs 2H1, 2H7 and 1D12 reduced α-syn p-Ser129 levels in DA neurons (N = 3-5 per genotype). (J) The schematic shows different DA neuronal clusters in an adult fly brain. Bar graphs are presented as mean ± SEM. One-way ANOVA, Dunnett’s multiple comparisons test. *p ≤ 0.05, ****p ≤ 0.0001 and ns = non-significant; Scale bar A-H’ = 100 μm.

Additionally, loss of PPM2 neurons was prevented by all sdAbs (2D10: p = 0.0075, 2H1: p = 0.0319, 2H7: p = 0.0035 and 1D12: p = 0.0075) except 2D8 (compare PPM2 region in Fig. S10D-H with C; see also J). These results indicate the effectiveness of sdAbs 2H1, 2H7, and 1D12 in reducing pathological levels of α-syn p-Ser129 and preventing neuronal loss in the vulnerable group of DA neurons.

### Anti-*α*-syn sdAbs increase the lifespan of synucleinopathy flies

The expression of α-syn has been shown to shorten the lifespan of *Drosophila* (*31*). Based on this, we also analyzed the lifespan of α-syn-expressing flies and whether it was affected by the anti-α-syn sdAbs. Compared to control flies, which typically survive to around 100 days, expression of α-syn and co-expression of Dv^sdAb^ limited to DA neurons, reduced lifespan to about 90 days [Fig. 6A; *TH>α-syn* (p = 0.0054)], *TH>α-syn+Dv^sdAb^* (p = 0.0073). This modest detrimental effect of a single copy of α-syn on survival agrees with a previously published report (*31*). In comparison to *TH>α-syn+Dv^sdAb^* flies, expression of sdAbs 2D8 and 1D12 did not affect lifespan. In contrast, expression of sdAbs 2D10 (p = 0.0038), 2H1 (p = 0.0039), and 2H7 (p = 0.0004) in the *TH>α-syn* background significantly increased fly lifespan (Fig. 6A). Additionally, we did not note any toxic effects of these sdAbs when driven exclusively in DA neurons (Fig. 6B) Overall, these findings indicate that anti-α-syn sdAbs not only improve function, prevent DA neuronal loss and lower levels of pathological α-syn p-Ser129 but also extend the lifespan of flies expressing α-syn in DA neurons.

**Fig. 6.**
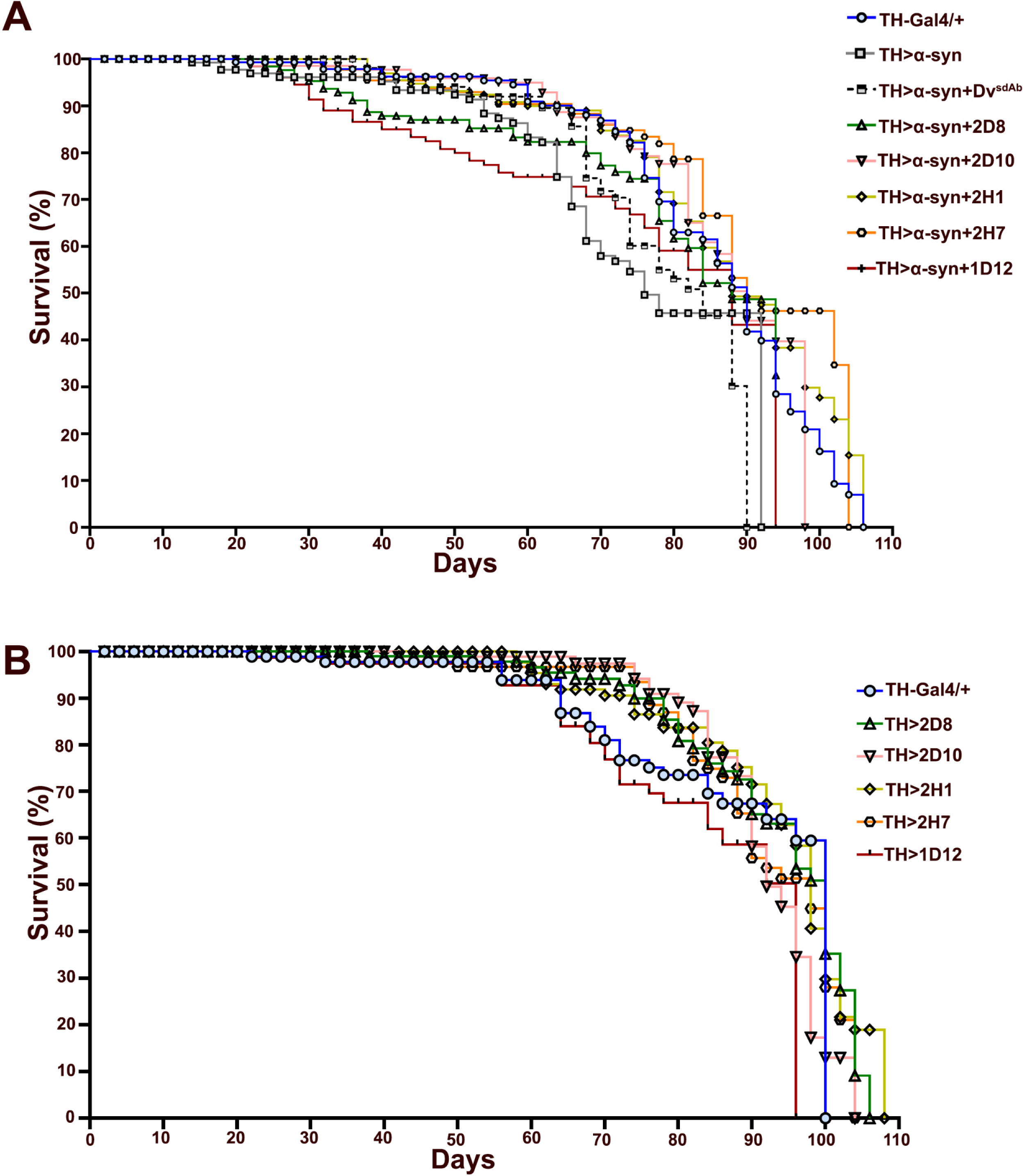
Anti-*α*-syn sdAbs improve the life span of synucleinopathy flies. (A) The expression of α-syn (p = 0.0054) without or with the negative sdAb control Dv^sdAb^ (p = 0.0073) in DA neurons reduced adult lifespan compared to control flies. The sdAb 2D8 (p = 0.19) and 1D12 (p = 0.8) did not extend lifespan compared with the negative control *TH>α-syn+Dv^sdAb^*. On the other hand, sdAbs 2D10 (p = 0.0038), 2H1 (p = 0.0039), and 2H7 (p = 0.0004) extended the life span compared to *TH>α-syn+Dv^sdAb^*flies, with the most effective 2H1 extending lifespan by 16 days. Genotypes and the number of flies analyzed per group: *TH-Gal4/+* (N = 65), *TH>α-syn* (N = 48), *TH>α-syn+Dv^sdAb^* (N = 38), *TH>α-syn+2D8* (N = 40), *TH>α-syn+2D10* (N = 37), *TH>α-syn+2H1* (N = 71), *TH>α-syn+2H7* (N = 48), and *TH>α-syn+1D12* (N = 50). (B) The effect of expressing the sdAbs alone (without α-syn) on the adult lifespan. The survival curves are similar between *TH-Gal4/+* controls and the different sdAbs, with no significant differences. Genotypes and the number of flies analyzed per group: *TH-Gal4/+* (N = 38), *TH>2D8* (N = 44), *TH>2D10* (N = 44), *TH>2H1* (N = 46), *TH>2H7* (N = 37) and *TH>1D12* (N = 26). Logrank (Mantel-Cox) test.

### Anti-*α*-syn sdAbs alleviate mitochondrial defects in flies

The association of α-syn with mitochondrial abnormalities has been extensively reported in various studies (*32–35*). Hence, we analyzed the efficacy of the anti-α-syn sdAbs in mitigating α-syn-induced mitochondrial defects. We thus utilized a *UAS-mito-Dsred* transgenic line to visualize mitochondrial morphology in different genotypes. Compared to control flies, which show thread-like mitochondrial morphology around the nucleus, pan-neuronal expression of α-syn and α-syn+Dv^sdAb^ resulted in a more punctate mitochondrial appearance (compare Fig. 7B-C to A; see also I). Compared with *Elav>α-syn+Dv^sdAb^*, mitochondrial appearance did not change in *Elav>α-syn+2D8* (Fig. 7D; see also I). However, expression of sdAbs 2D10 (p = 0.0009), 2H1 (p <0.0001), 2H7 (p <0.0001), and 1D12 (p <0.0001) in the *Elav>α-syn* background changed the mitochondrial morphology from punctate to thread-like (Fig. 7E-H; see also I). We also analyzed the efficacy of these sdAbs on superoxide levels in live mitochondria utilizing MitoSOX. Compared to control, pan-neuronal expression of α-syn or α-syn+Dv^sdAb^ increased the superoxide levels (compare Fig. S11B with A; see also I, J). Except for 2D8, co-expression of anti-α-syn sdAbs 2D10 (p = 0.0002), 2H1 (p = 0.0276), 2H7 (p = 0.0088), and 1D12 (p <0.0001) reduced the superoxide levels (compare Fig. S11C with D to H; see also J).

**Fig. 7.**
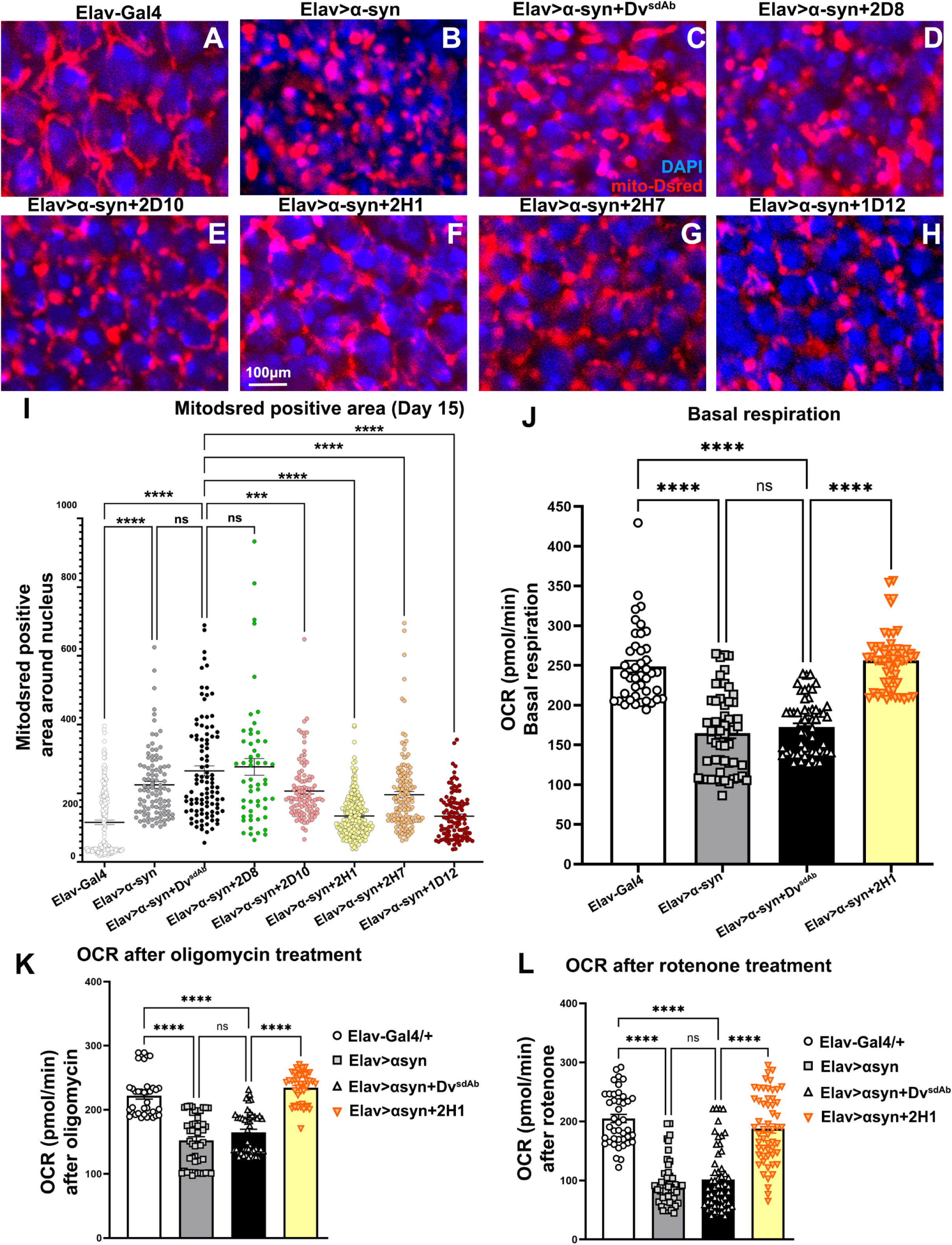
Pan neuronal expression of anti-*α*-syn sdAbs mitigates mitochondrial dysfunctions in *α*-syn expressing flies. (A-H) Confocal images of 15-day-old female adult brains expressing mito-Dsred (Dsred with a mitochondrial targeting sequence) and DAPI. (I) Quantification of mito-Dsred positive area around the nucleus in different age-matched genotypes shows that all sdAbs except 2D8 reduced mito-Dsred positive area, which indicates a change in mitochondrial morphology from punctate to thread-like in α-syn expressing flies (N=5-8 per genotype). (J-L) The bar graph shows basal respiration OCR levels (J), OCR after oligomycin (K), and OCR after rotenone administration (L) in different genotypes. 2H1 restored OCR to near-control levels in synucleinopathy flies (N = 6-8 per genotype). Scatter plot graphs are presented as mean ± SEM. One-way ANOVA, Tukey’s multiple comparisons test. ***p ≤ 0.001, ****p ≤ 0.0001 and ns = non-significant; Scale bar A-H = 100 μm.

Collectively, of the different sdAbs, 2H1 consistently rescued every measured aspect of α-syn pathology. Therefore, for subsequent analysis, we only assessed the efficacy of sdAb 2H1.

After analysing mitochondrial morphology, we investigated mitochondrial function by Seahorse analysis to assess changes in oxygen consumption rate (OCR). We measured OCR changes by injecting oligomycin and rotenone, which inhibit distinct mitochondrial complexes. Compared with control, pan-neuronal expression of α-syn and α-syn+Dv^sdAb^ reduced basal respiration (Fig. 7J). Interestingly, pan-neuronal expression of 2H1 in synucleinopathy flies restored basal OCR to near-control levels (p <0.0001; Fig. 7J). A similar pattern of OCR changes was observed after injecting oligomycin (Elav>α-syn+Dv^sdAb^ vs Elav>α-syn+2H1: p <0.0001; Fig. 7K) and rotenone (Elav>α-syn+Dv^sdAb^ vs Elav>α-syn+2H1: p <0.0001; Fig. 7L). A complete curve of the Seahorse analysis is also shown in Fig. S11K. This indicates that pan-neuronal expression of anti-α-syn sdAbs not only restricts changes in mitochondrial morphology but also reverses reduced OCR to near-normal levels.

### Epitope mapping of anti-*α*-syn sdAbs against peptides spanning *α*-synuclein and binding affinity

To clarify the variable efficacy of anti-α-syn sdAbs, we determined their α-syn epitopes and binding affinities. To assess the binding domains of anti-α-syn sdAbs, we performed a dot blot assay using 10 to 15 amino acid peptides covering overlapping regions of the whole 140–amino acid α-syn. (Fig. 8A). The sequences of each α-syn peptide are listed in Table S1. We used bovine serum albumin (BSA) and the soluble fraction [low-speed supernatant (LSS)] from a tg M83 mouse brain expressing A53T α-syn as negative and positive controls, respectively. We previously reported that sdAb 2D8 reacts primarily with peptides 8 and 9 [α-syn 71 to 95; Fig. 8B (*15*)], whereas 2D10 binds strongly to peptide 9 [α-syn 81 to 95; Fig. 8C; (*15*)]. For the other anti-α-syn sdAbs, 2H1 bound strongly to peptides 7, 8, and 9 (α-syn 61 to 95); 2H7 bound to peptide 8 (α-syn 71 to 85), while 1D12 bound to peptide 7 (α-syn 61 to 75; Fig. 8D-F; see also Fig. S12A-C). These epitopes comprise the hydrophobic domain (amino acids 61-95) of α-syn, which is responsible for its aggregation (*36, 37*). In that region, the amino acid stretch GAVV (residues 68–71) is essential for nucleation and aggregation into prion-like fibrils, whereas residues 79–95 are important for stabilizing these fibrils and facilitating seeding competency (*37*).

**Fig. 8.**
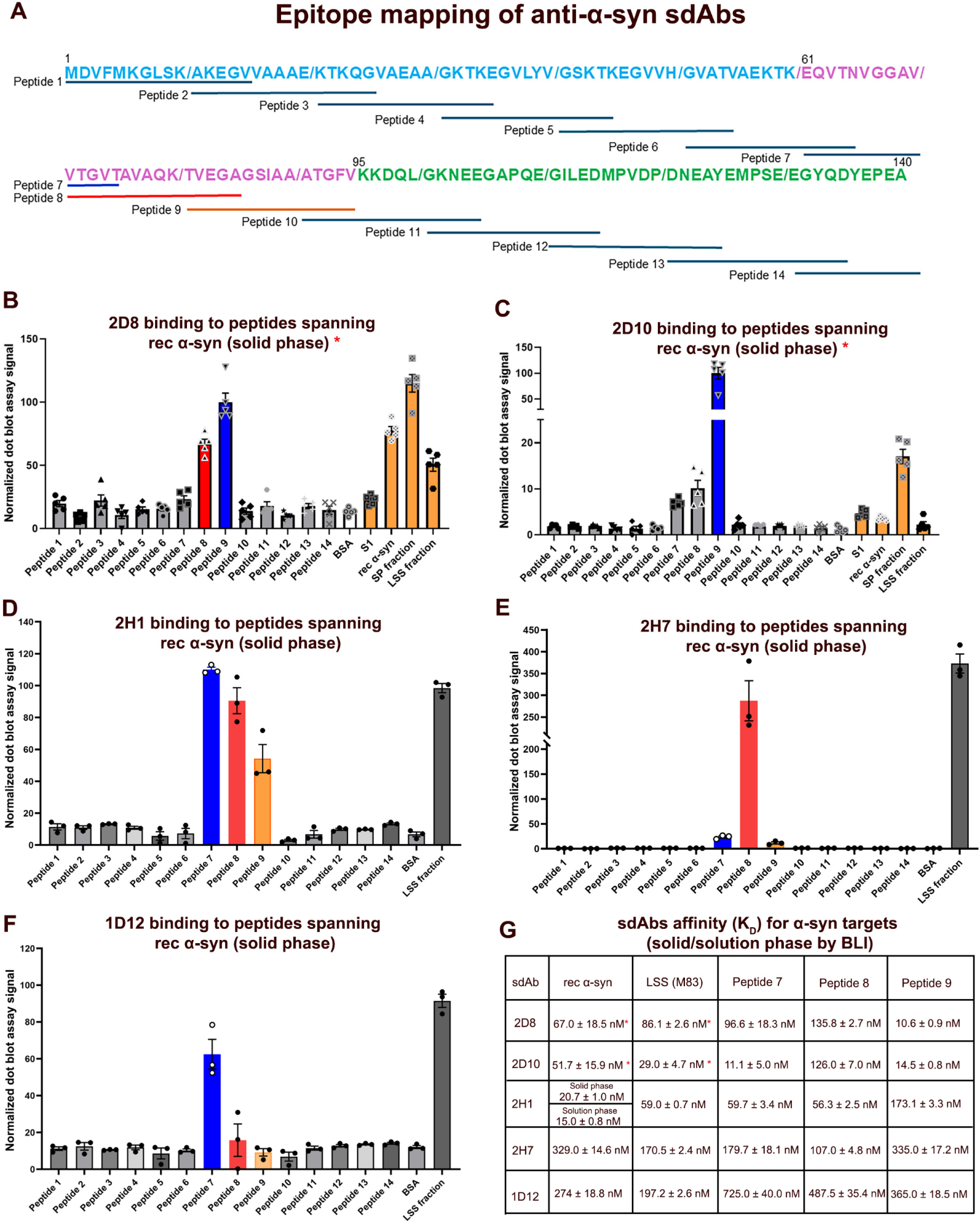
Epitope mapping and binding affinities of all anti-*α*-syn sdAbs by dot blot and BLI binding affinity assay with *α*-syn targets. (A) The schematic depicts a peptide library of α-syn, with each peptide overlapping the next by 5 amino acids. Blue indicates the N-terminal amphipathic repeat region (amino acids 1 to 60, membrane-binding domain), pink highlights the hydrophobic domain (amino acids 61 to 94, which promotes aggregation), and green marks the C-terminal acidic domain (amino acids 95 to 140). (B-F) Quantification of the dot blot assay results for 2D8 (B), 2D10 (C), 2H1 (D), 2H7 (E), and 1D12 (F). 2D8 reacted mainly with peptides 8-9, 2D10 with peptide 8, 2H1 with peptides 7-9, 2H7 with peptide 8, and 1D12 with peptide 7. Positive control: LSS soluble fraction from a tg-M83 mouse brain; negative control: BSA. Bar graphs are presented as mean normalized signal ± SEM from three replicates. (G) sdAb affinity (K_D_) for α-syn and its related peptides targets in the solid and solution phase. * Some data for 2D8 and 2D10 have been previously published (*15*).

Subsequently, we analyzed binding affinities in the solution phase using a biolayer interferometry (BLI) assay. First, we assessed the affinities of sdAbs for the rec α-syn and the soluble LSS fraction of tg M83 mouse brain homogenate. We previously reported that 2D8 and 2D10 have high affinity for rec α-syn (2D8: *K_D_* = 67.0 ± 18.5 nM; 2D10: *K_D_* = 51.7 ± 15.9 nM) and LSS [2D8: *K_D_ =* 86.1 ± 2.6 nM; 2D10: *K_D_ =* 29.0 ± 4.7 nM; Fig 8G; (*15*)]. For the other sdAbs, 2H1 had high affinity (rec α-syn: *K_D_* = 15.0 ± 0.8 nM; LSS: *K_D_ =* 59.0 ± 0.7 nM; Fig. S12D, H and Fig. 8G), while 2H7 and 1D12 had moderate to low affinity for rec α-syn (2H7: *K_D_* = 329.0 ± 14.6 nM; 1D12: *K_D_* = 274.0 ± 18.8 nM; Fig. S12F-G and Fig. 8G) and LSS (2H7: *K_D_ =* 170.5 ± 2.4 nM; 1D12: *K_D_ =* 197.2 ± 2.6 nM; Fig. S12I-J and Fig. 8G).

Next, we analyzed the affinities of these sdAbs for peptides 7, 8, and 9. 2D8 had the highest affinity for peptide 9 (*K_D_* = 10.6 ± 0.9 nM; Fig. 8G and Fig. S13C), followed by moderate binding to peptide 7 (*K_D_* = 96.6 ± 18.3 nM; Fig. 8G and Fig. S13A) and the lowest interaction with peptide 8 (*K_D_* = 135.8 ± 2.7 nM; Fig. 8G and Fig. S13B). 2D10 had the highest affinity for peptide 7 (*K_D_* = 11.1 ± 5.0 nM; Fig. 8G and Fig. S13D) and 9 (*K_D_* = 14.5 ± 0.8 nM; Fig. 8G and Fig. S13F), and the lowest for peptide 8 (*K_D_* = 126.0 ± 7.0 nM; Fig. 8G and Fig. S13E). 2H1 had the highest affinity for peptide 7 (*K_D_* = 59.7 ± 3.4 nM; Fig. 8G and Fig. S13G) and peptide 8 (*K_D_* = 56.3 ± 2.5 nM; Fig. 8G and Fig. S13H), and the lowest for peptide 9 (*K_D_* = 173.1 ± 3.3 nM; Fig. 8G and Fig. S13I). 2H7 had a moderate affinity for peptide 8 (*K_D_* = 107.0 ± 4.8 nM; Fig. 8G and Fig. S13K) and low for peptide 7 (*K_D_* = 179.7 ± 18.1 nM; Fig. 8G and Fig. S13J) and peptide 9 (*K_D_* = 335.0 ± 17.2 nM; Fig. 8G and Fig. S13L). On the other hand, 1D12 had the lowest affinity for all three peptides (peptide 7: *K_D_* = 725.0 ± 40.0 nM; peptide 8: *K_D_* = 487.5 ± 35.4 nM, and peptide 9: *K_D_* =365.0 ± 18.5 nM; Fig. 8G and Fig. S13M-O). Of the sdAbs, 2H1 had the highest affinity for rec α-syn and high affinity for LSS and for peptides 7 and 8 (Fig. 8G). In view of its consistent rescue efficacy in the synucleinopathy flies, we also evaluated the affinity of 2H1 for rec α-syn in the solid-phase, to which it showed high affinity (*K_D_* = 20.7 ± 1.06 nM; Fig. 8G and Fig. S12E). Notably, all the anti-α-syn sdAbs examined here were derived from solid-phase panning (*15*). Collectively, these affinity studies (summarized in Fig. 8G) likely explain 2H1’s prolonged and consistent rescue efficacy across all performed assays in the flies.

### TurboID-based identification of 2H1 interacting proteins in neurons expressing *α*-synuclein

Among all the examined anti-α-syn sdAbs, 2H1 was consistently effective in inhibiting all outcome measures of α-syn-mediated pathology. To investigate how 2H1 mediates rescue, we profiled the in vivo interactors to 2H1 and α-syn through proximal ligation by fusing turboID-V5 to our proteins of interest (Fig. 9A). TurboID is a bioengineered enzyme that labels proteins within approximately 10 nm via proximity-dependent biotinylation (*38*). We created transgenic lines *UAS-2H1-turboID* and *UAS-Dv^sdAb^-turboID*, with turboID fused to the C-terminus with a V5 tag (Fig. 9B). The *UAS-Dv^sdAb^-turboID* serves as a negative control; its size is comparable to 2H1 (∼15 kDa; Fig. 9B), and it does not bind to α-syn, which allows for identifying non-specific interactions.

**Fig. 9.**
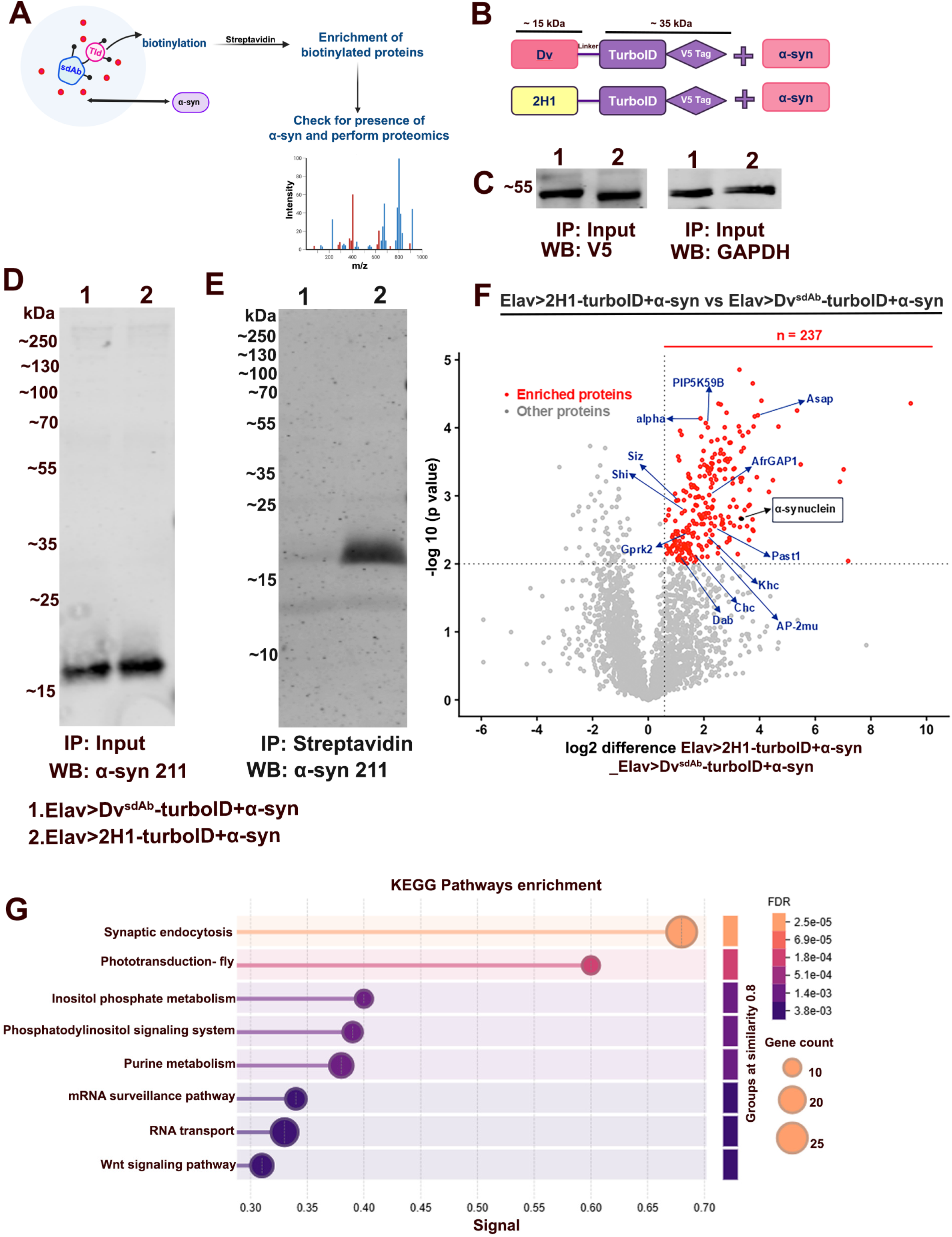
TurboID-tagged sdAb 2H1 shows effective in vivo association with *α*-syn and enrichment of endocytosis-related factors in the *Drosophila* model of synucleinopathies. (A) The schematic represents the strategy for enriching proteins biotinylated by 2H1 turboID and their analysis by LC-MS in α-syn expressing flies. (B) Control Dv^sdAb^ turboID and 2H1 turboID constructs were designed with turboID tags at the C-termini of the respective proteins. (C) Western blot analysis with immunoprecipitation (IP) input confirmed the expression of the V5 tag in different turboID constructs co-expressed with α-syn. (D) Western blot analysis with IP input confirmed the presence of α-syn in *Elav>*Dv^sdAb^*-turboID+α-syn* and *Elav>2H1-turboID+α-syn.* (E) Co-IP using streptavidin pulldown followed by detection of α-syn antibody. α-syn was detected in *Elav>2H1-turboID+α-syn* (lane 2) but not in *Elav>*Dv^sdAb^*-turboID+α-syn* (lane 1), confirming in vivo association of 2H1 to α-syn. (F) Volcano plot shows proteins differentially biotinylated in *Elav>2H1-turboID+α-syn* samples versus *Elav>Dv^sdAb^-turboID+α-syn* fly samples from on-bead digestion experiments. Two hundred thirty-seven proteins were significantly biotinylated (p < 0.01 and a fold change of >1.5) by 2H1-turboID as compared to Dv^sdAb^-turboID in α-syn expressing flies. The highlighted proteins are related to the endocytosis pathway (in blue). Also highlighted is α-syn (in black) that was preferentially biotinylated in *Elav>2H1-turboID+α-syn* flies. (G) KEGG pathways enrichment analysis for proteins preferentially biotinylated in 2H1-turboID+α-syn as compared to Dv^sdAb^-turboID+α-syn flies.

We expressed *UAS-2H1-turboID* or *UAS-Dv^sdAb^-turboID* pan-neuronally in *Elav>α-syn* flies and verified α-syn expression with the syn211 antibody. Input samples from both *Elav>UAS-2H1-turboID+α-syn* and *Elav>UAS-Dv^sdAb^-turboID+α-syn* displayed α-syn at around 15 kDa with equal loading and notable expression of V5 tag (Fig. 9C, D). After confirming α-syn expression in input samples, we enriched biotinylated proteins using streptavidin beads and probed for total α-syn using the syn211 antibody. Compared to the negative control *Elav>UAS-Dv^sdAb^ -turboID+α-syn* that did not show notable binding with α-syn (lane 1, Fig. 9E; also compare Fig. 9E with D), *Elav>UAS-2H1-turboID+α-syn* displayed α-syn at about 15 kDa, confirming in vivo binding of 2H1 to α-syn (compare lane 2 with lane 1, Fig. 9E).

After confirming the in vivo interaction between 2H1 and α-syn, we expanded our analysis to profile the 2H1-turboID interactome when co-expressed pan-neuronally with α-syn. The proteome of samples co-expressing 2H1-turboID+α-syn was compared with the negative control expressing Dv^sdAb^-turboID+α-syn. Using a significance threshold of p < 0.01 and a fold change of >1.5, we identified 237 proteins significantly biotinylated by 2H1-turboID as compared to Dv^sdAb^-turboID (Fig. 9F). We also noted the presence of α-syn, which confirms that 2H1 binds to it in vivo (Fig. 9F).

A complete list of significantly enriched interactors is shown in Supplementary Table S2. Applying String software to enriched proteins (filtered at FDR ≤ 0.05) shows that the most enriched GO terms, based on the highest Combined Score indicating the strongest enrichment, are mainly related to synapse and vesicle-trafficking pathways. KEGG pathway showed enrichment for endocytosis, followed by phototransduction, inositol phosphate metabolism, phosphatidylinositol signaling system, purine metabolism, mRNA surveillance pathway, RNA transport, and Wnt signaling pathway (Fig. 9G). Furthermore, categorization of enriched GO terms based on their biological, molecular, and cellular component is shown in Fig. S14A-C. All these results indicate that enriched proteins are involved in the synaptic vesicle trafficking pathway at the nerve terminal. Of note, independent proximal ligation screens with α-syn in cultured mammalian cells have also reported α-syn’s association with synaptic and endocytic proteins (*39, 40*). Moreover, α-syn aggregates trap vital synaptic proteins like VAMP2/synaptobrevin-2 and SNAP25, causing synaptic dysfunction in PD and other synucleinopathies (*41–43*). Therefore, many proteins specifically biotinylated by 2H1-turboID could be α-syn-associated factors, which we plan to explore in future studies.

## Discussion

Synucleinopathies such as PD, DLB, and MSA involve the accumulation of α-syn aggregates, including Lewy bodies and glial inclusions (*1*). Immunotherapies targeting α-syn are promising (*44–47*), but IgG antibodies face challenges in crossing the BBB due to their large size (∼150 kDa). In contrast, sdAbs (∼15 kDa) have much better brain penetration, can bind to cryptic epitopes, and are ideal for gene therapy because of their single unit. We developed sdAbs against human α-syn that target its hydrophobic domain, unlike the IgGs in clinical trials that target its N- or C-terminus.

In this study, we evaluated the therapeutic potential of different anti-α-syn sdAbs first by testing them in vitro in primary tg A53T α-syn mouse neurons incubated with α-syn enriched fraction from human DLB brain, and then in vivo in a *Drosophila* model of synucleinopathy that expresses codon-optimized human α-syn and exhibits classic symptoms of motor defects and DA neuronal loss (*20, 26*). In culture, all sdAbs except one (1D12) prevented α-syn aggregation while some prevented its toxicity (Fig. 1). Additionally, in vivo data indicated that all the sdAbs alleviated motor deficits in females at 15 days post-eclosion, while only 2H1 remained effective at day 25, compared with the negative control sdAb Dv^sdAb^ (Fig. 2). However, we noted that these anti-α-syn sdAbs were ineffective in attenuating motor impairments in males (Fig. S5). This may be due to dosage compensation in males (*48, 49*), which leads to higher expression of Gal4-targeted genes, thereby increasing α-syn toxicity and reducing the efficacy of anti-α-syn sdAbs, as supported by more severe locomotor deficits in the males compared to the females (compare Fig. 2 with S5). Further, we examined the potency of these sdAbs in inhibiting α-syn-mediated DA neuronal loss (*27*). We noted that the loss of PPL1 was prevented by pan-neuronal co-expression of sdAbs 2D8, 2D10, and 2H1, whereas sdAbs 2H7 and 1D12 were ineffective. Similarly, degeneration of the PPM2 cluster was prevented by sdAbs 2D10, 2H1, and 1D12, while sdAbs 2D8 and 2H7 were ineffective (Fig. 3). Presumably related to its effect in the climbing assay, 2H1 prevented loss of the PPL1 neuronal population through 25 days post-eclosion (Fig. S7). These results indicate that of the examined sdAbs, 2H1 was most effective at preventing both motor impairments and DA neuron loss during aging.

In PD, α-syn undergoes numerous post-translational modifications (PTMs), including nitration, phosphorylation (*50*), and modifications regulated by DA (*51*). In PD, Lewy bodies predominantly contain α-syn phosphorylated at Ser129 (*6, 52*). Additionally, soluble, non-fibrillar α-syn in PD tissues is phosphorylated at Ser129 (*8*), suggesting that this modification may influence its aggregation and oligomerization, implicating a connection to PD pathology. α-syn phosphorylation at Ser129 is strongly associated with pathology (*6–8, 30*), therefore, we evaluated the effectiveness of these sdAbs in controlling pathogenic α-syn phosphorylation levels. We noted that sdAbs 2H1, 2H7, and 1D12 reduced α-syn p-Ser129 but not total α-syn levels in 15-day-old synucleinopathy flies (Fig. 4). This highlights the specificity of these sdAbs in targeting pathological α-syn levels. Additionally, we examined the efficacy of sdAbs in reducing this epitope in DA neurons undergoing degeneration, which is prominent in different synucleinopathies (*22*). We noted a similar efficacy pattern, wherein sdAbs 2H1, 2H7, and 1D12 reduced α-syn p-Ser129 levels (Fig. 5). This highlights the potency of these sdAbs in clearing toxic α-syn in all neurons, including vulnerable DA neurons. We also noted that sdAbs 2D10, 2H1, and 2H7 increased fly lifespan when α-syn was expressed exclusively in DA neurons (Fig. 6). This reiterates the overall ability of these sdAbs to eliminate the toxic effects of pathogenic α-syn.

Numerous studies have enhanced our knowledge of how α-syn is linked to mitochondrial abnormalities. For instance, oligomeric α-syn alters complex I-regulated respiration and oxidizes the ATP synthase β subunit, leading to mitochondrial swelling and cell death (*12*). In addition, α-syn p-Ser129 aggregates interact with mitochondria and impair their cellular respiration (*11*). In flies, α-syn expression causes mitochondrial dysfunction by mislocalizing Drp1, a key mitochondrial fission protein (*35*). Considering this evidence, we felt compelled to test whether our sdAbs could rescue mitochondrial defects. We observed that α-syn expression altered mitochondrial morphology, which was attenuated by sdAbs 2D10, 2H1, 2H7, and 1D12. Additionally, we performed Seahorse analysis and found that pan-neuronal expression of human α-syn reduced OCR, which was prevented by 2H1 (Fig. 7). Therefore, our results are consistent with the published literature regarding the toxic effects of α-syn on mitochondria, which was mitigated by the anti-α-syn sdAbs.

To gain insight into potential reasons for differences in sdAbs’ efficacy, we determined their α-syn epitopes and examined their binding affinities for the immunogen, full-length rec α-syn (140 aa), and overlapping peptides spanning the hydrophobic region of α-syn in the solid and solution phases, respectively. In the solid phase, we noted that anti-α-syn sdAbs targeted the central hydrophobic region (aa 61-95), but their binding profile within this region differed. 2D8 bound to aa 71-95 (peptides 8 and 9), 2D10 to aa 81-95 (peptide 9), 2H1 to aa 61-95 (peptides 7, 8, and 9), 2H7 to aa 71-85 (peptide 8), and 1D12 to aa 61-75 (peptide 7). Only 2H1 bound to the entire hydrophobic region that governs α-syn aggregation [Fig. 8; (*36, 37*)], which likely accounts for its prolonged and consistent rescue efficacy across assays. Afterward, we noted that in the solution phase, 2D10 and 2H1 had the highest affinity to LSS and rec α-syn, respectively. Further, 2D10, 2H1, and 2D8 had the highest affinity to peptides 7, 8, and 9, respectively (Fig. 8G). Collectively, the ability of 2H1 to bind to the entire hydrophobic region in the solid phase (Fig. 8D) and its high binding affinity for the major hydrophobic region and rec α-syn in the solution phase (Fig. 8G) might explain its prolonged and consistent in vivo rescue efficacy.

To further validate 2H1’s function, we developed turboID constructs to identify proteins that interact with 2H1 sdAb in α-syn expressing flies. Streptavidin-mediated enrichment of biotinylated proteins from 2H1-turboID+α-syn flies confirmed an in vivo interaction between 2H1 and α-syn, but not with the negative control Dv^sdAb^-turboID+α-syn (Fig. 9). Subsequently, proteomics validated 2H1’s interaction with α-syn. GO term pathway analysis of 2H1-turboID+α-syn-enriched proteins showed those involved in endocytosis and synaptic vesicle cycle pathways (Fig. 9G; Fig. S14A-C). We note that independent proximity ligation screens with α-syn also identified proteins involved in synaptic and endocytic function (*39, 40*). Moreover, α-syn sequesters synaptic proteins such as SNAP25 and VAMP2 and thereby impairs synaptic function (*43*). These findings indicate that 2H1 binds α-syn and associates with its interactome in vivo. Proteomic analysis of 2H1-turboID+α-syn mediated biotinylated proteins showed enrichment of genes Rpt2, Rpn1, and Prosalpha1, which are key components of the proteasome subunit (*53*). This might explain the sdAb-mediated reduction in α-syn p-Ser129 levels. Additionally, enrichment of genes DA 2-like receptor (D2R; activated by DA) and pale (ple; tyrosine hydroxylase) with 2H1-turboID+α-syn complex supports our results demonstrating sdAb-mediated prevention of DA neuron loss. Furthermore, genes such as Vps13D, Aralar1, Pgam5, and Drp1, which are key regulators of mitochondrial dynamics (*54–57*), are also enriched in our interactome. This supports our results, which demonstrate that sdAb alleviates α-syn-mediated mitochondrial defects.

We previously demonstrated that anti-syn sdAbs 2D8 and 2D10 enable specific, non-invasive visualization of α-syn in vivo, highlighting their potential for diagnostic imaging. These monoclonal sdAbs were readily detected in neurons, where they bound α-syn and were localized to early endosomes, late endosomes/lysosomes, and autophagosomes (*15*). To enhance the therapeutic potential of 2D8, we conducted a follow-up study in which we fused a protein degrader to the sdAb to promote ubiquitination and proteasomal degradation of α-syn. This approach enhanced proteasomal degradation of α-syn, complementing the endogenous lysosomal degradation pathway. This modified sdAb improved α-syn clearance in primary cell cultures and in a mouse model of synucleinopathy (*16*). These studies highlight the ability of unmodified and modified sdAbs to target misfolded/aggregated α-syn via the proteasomal/lysosomal pathways, which supports findings from the present study as well.

In addition to our sdAbs targeting the hydrophobic α-syn region, several anti-α-syn sdAbs have been reported in the literature that mostly target the N- or C-terminus of α-syn and have primarily been characterized in vitro (*58–69*). A few have been examined in vivo (*70, 71*). Two sdAbs injected intracerebrally as AAV constructs showed varying efficacy in different readouts in a rat model based on an AAV-α-syn brain injection (*70*). Another synthetic sdAb administered intraventricularly in an AAV construct impeded the spreading of brain injected fibrillar α-syn in mice (*71*). In contrast, our sdAbs targeting the hydrophobic region are not only efficacious in culture but also in mouse and fly models, and we have now identified one particular sdAb, 2H1, that is substantially more efficacious than 2D10 and 2D8, the two that we have described previously (*15, 16*).

Taken together, our findings detail the in vivo efficacy of different anti-α-syn sdAbs in preventing α-syn-mediated death as well as functional, cellular and molecular defects in *Drosophila* synucleinopathy models. sdAb 2H1 binding to the entire hydrophobic α-syn region provides the highest efficacy in preventing synucleinopathy, which supports its clinical development. Lastly, we provide insight into its mechanism of action, which likely takes place in the synapse, where α-syn is highly concentrated.

## Materials and methods

### Synuclein fractionation

Frozen brain tissue from a patient with extensive cortical α-syn deposits, indicative of Lewy Body Dementia, was obtained from National Disease Research Interchange (Philadelphia, PA). Following previously published protocol, the tissue was homogenized in TBS (Tris-Buffered Saline) and centrifuged at 3000 x g for 20 min to generate the first supernatant S1 fraction (*16*). This fraction contains the total soluble synuclein species and was found to be the most toxic in culture.

### Neuronal culture

Primary neuronal cultures were prepared from pups at postnatal day 0 using methods previously described (*72–74*). For these experiments we utilized mice from the M83 line which expresses human α-syn containing the familial A53T mutation [JAX #004479, (*75*)]. The cortex and hippocampus were collected, washed in modified Hank’s Balanced Salt Solution (HBSS), and then digested for 30 min at 37°C with trypsin. The tissue was then washed again and manually dissociated. Cells were added to 24 well culture plates and incubated in plating media for 24 h, after which it was replaced with Neurobasal media.

### Synuclein and sdAb treatment

After allowing the cells to recover from plating, cultures were exposed to 10 µg/ml of S1 extract and 1 μg/ml of the sdAbs in one of two dosing paradigms. In the α-syn → sdAb method, the α-syn was added first and allowed to incubate in culture for 24 h, after which the remaining α-syn in the media was washed off, and the sdAb was then added. In this paradigm, the pathological α-syn is internalized, and the sdAb binds to it inside the neuron. On the other hand, in the α-syn + sdAb paradigm the S1 and sdAbs were added to the media together, which simulates interaction between α-syn and the sdAbs in the extracellular space.

### Drosophila stocks

In the present study, *Drosophila* stocks were reared in standard cornmeal/agar/yeast media at 25°C on a 12h:12h light: dark cycle, according to standard protocol. For modeling synucleinopathies in *Drosophila,* we used previously described codon-optimized human α-syn transgenic lines (*20*). *Elav-Gal4/Y; UAS-SNCA/CyoGal80*; *+/+* was generously provided by Professor Juan Botas (*21*). To study the efficacy of anti-α-syn sdAbs, each sdAb transgene, tagged with myc and his at the C-terminus, was subcloned into the pUAST-attB plasmid and targeted for insertion at a specific locus on the 2nd chromosome (51C) to create a stable transgenic line (BestGene Inc.) for sdAbs: *w; UAS-2D8; +/+*, *w; UAS-2D10;+/+*, *w; UAS-2H1;+/+*, *w; UAS-2H7;+/+*, and *w; UAS-1D12;+/+*, wherein UAS is an upstream activator sequence where Gal4 binds to regulate expression of a downstream gene (*76*). For our manuscript, we used the previously described negative control of UAS-Dv^sdAb^ (*17*). We have also generated transgenic lines: *w; UAS-Dv^sdAb^-turboID; +/+*, and *w; UAS-2H1-turboID; +/+* with turboID along with a V5 tag at their C-terminal. We subcloned the respective constructs into the pUAST-attB plasmid, which was inserted at a specific locus on the 2nd chromosome (51C) to create stable transgenic lines (BestGene Inc.). Other transgenic lines used in this study: elav^c155^-GAL4 (drives expression in all neurons; BL #458), W^1118^ (BL #5905), UAS-mito-Dsred (BL #93056), TH-Gal4 (drives expression in DA neurons; BL #8848), were obtained from the Bloomington Drosophila Stock Center, Bloomington, Indiana, USA.

### Climbing and survival assays

To perform the climbing assay, 1-day-old flies were collected and transferred to fresh food vials, with up to 10 adults per vial. Female and male flies from each genotype were then transferred into separate empty test tubes marked at 8 cm. After a 1-minute rest, the test tube was gently tapped, a 10-second timer was started, and the number of flies crossing the mark was recorded. This process was repeated three times for each set of vials, with 1-minute intervals between readings, and was recorded with an iPhone camera. The assay was conducted up to 25 days after eclosion, and the percentage of flies crossing the mark was calculated and visualized using GraphPad Prism 10.0.2. For the female analysis, 100-194 flies per genotype were tested, divided into sets of 10-22 flies. For male analysis, 42-97 flies per genotype were tested, divided into sets of 5-10 flies. The average of each set was calculated and plotted as an individual data point.

For the survival assay, adult female flies of the respective genotypes were collected and kept in fresh food vials, with up to 10 flies per vial. The number of surviving flies was counted and recorded daily. Every other day, flies were moved to new food vials. The log-rank (Mantel-Cox) test was employed to assess differences between genotypes using GraphPad Prism 10.0.2.

### Immunostaining

For immunostaining, adult *Drosophila* brains were dissected, fixed in 4% paraformaldehyde [PFA, diluted in phosphate-buffered saline (PBS)], and washed with PBS-T (PBS containing 0.5% Triton X-100, pH 7.4). Samples were then blocked with 5% normal goat serum in PBS-T and incubated with the specified primary antibody overnight at 4°C. Primary antibodies used for immunostaining were rabbit anti-tyrosine hydroxylase (1:500, Millipore Sigma, Cat #AB152) and rabbit anti-α-syn p-Ser129 (1:1000, Invitrogen, Cat #PA1-4686), and mouse anti-tyrosine hydroxylase (1:1000, Immunostar, Cat #22941). Tissues were incubated with their respective secondary antibody (Alexa Fluor, 1:500, Invitrogen) for 2 h at room temperature. Subsequently, the samples were washed three times with PBS-T and once with 1X PBS. Afterward, they were mounted using Vectashield antifade mounting media with DAPI (Vector Laboratories) and imaged with a Zeiss LSM700 confocal microscope.

### Protein extraction and western blotting

For protein extraction, adult female flies were decapitated and homogenized using RIPA buffer [0.5% Na-deoxycholate, 1% Triton X-100, 0.05 M Tris-Cl (pH-8.0), 0.02% Na-Azide, 0.1% SDS, 0.15 M NaCl], along with a phosphatase inhibitor cocktail (Pierce; NaF, Na_3_VO_4_, Na_4_P_2_O_7,_ C_3_H_7_Na_2_O_6_P) and a protease inhibitor cocktail (cOmplete, Roche). The homogenate was then centrifuged at 12,000 rpm for 10 min to pellet cellular debris, and the supernatant was collected and stored at -80°C until further use.

The total protein content was measured using a Pierce BCA Protein Assay Kit, and an equal amount was loaded onto a 12% or 15% SDS-PAGE gel. Following separation, the protein bands were transferred onto a polyvinylidene difluoride (PVDF) or nitrocellulose membrane (Millipore, USA) and blocked in Superblock (Thermo Fisher Scientific) or 5% non-fat dry milk prepared in TBS-T buffer (1× Tris-buffered saline with 0.1% Tween 20) for 1 to 2 h at room temperature. The lysates were then incubated with the specific primary antibodies at 4°C overnight. Primary antibodies utilized for western blotting were goat IgG anti-sdAb (VHH) conjugated with peroxidase (1:1000, Jackson ImmunoResearch, Cat #128-035-232), mouse anti-GAPDH (1:3000, Santa Cruz, Cat #365062), rabbit anti-α-syn p-Ser129 (1:1000, Invitrogen, Cat # PA1-4686), mouse anti–α-syn 211 (1:1000, Invitrogen, Cat #32-8100), rat anti-Elav (7E8A10, 1:1000, DSHB), and mouse anti-V5 (1:1000, Invitrogen, Cat #R960-25). The next day, the blot was adequately washed with 1X TBS-T (1X TBS with 0.1% Tween 20) and incubated with IRDye 800CW or IRDye 680RD secondary antibodies (1:10,000; LI-COR Biosciences) for 2 h at room temperature. Then, the blot was washed with 1X TBS-T and developed using the Odyssey XF Imaging System. All immunoblots were quantified using LI-COR Image Studio Lite 5.2. To detect sdAbs (VHH), the chemiluminescent signal was visualized using a Fuji LAS-4000. GraphPad Prism 10.0.2 was used for plotting relative signal intensity.

### Quantitative real-time PCR

RNA was extracted from 5-day-old adult fly heads using TRIzol reagent (Invitrogen, Cat # 15596026). cDNA synthesis was performed with the Verso cDNA synthesis kit (Thermo Scientific, Cat # AB1453A) using 1 μg of total RNA. 10 ng cDNA was used for qPCR, which employed Power SYBR Green mix (Applied Biosystems, Cat #4367659) and gene-specific primers listed below. The qPCR cycling conditions were 95°C for 5 min, followed by 40 cycles at 95°C for 10 s and 60°C for 45 s, ending with a melting curve. Rp49 served as the internal reference gene. Three independent biological replicates, each with three technical replicates per genotype, were analyzed. RP49 forward: 5’ GGTTTCCGGCAAGCTTCAA 3’, RP49 reverse: 5’ TGTTGTCGATACCCTTGGGC 3’; sdAb forward: 5’ ACTGCAACTACTGTGAAAT, sdAb reverse: 5’ ACTGCAACTACTGTGAAATCTGCCA 3’.

### Enrichment of biotinylated proteins by streptavidin pulldown

Adult female flies were collected and aged for 10 days on regular fly food. Flies were then depleted of biotin for ∼3-4 days by feeding on a 1% sucrose:1.5% agar solution under aseptic conditions. After depletion, flies were fed 100 µM biotin reconstituted in 1% sucrose:1.5% agar food. We depleted biotin to reduce the endogenous biotinylated proteins. Fifteen-day-old adult female flies (20 per biological replicate) were decapitated and homogenized in RIPA buffer supplemented with a phosphatase inhibitor cocktail (Pierce; NaF, Na_3_VO_4_, Na_4_P_2_O_7,_ C_3_H_7_Na_2_O_6_P) and a protease inhibitor cocktail (cOmplete, Roche). The homogenate was centrifuged at 12,000 rpm for 10 min to remove cellular debris, and the supernatant was collected and stored at -80°C until further analysis. Protein concentration was measured in each sample with a BCA protein assay. For pull-down of biotinylated proteins, streptavidin magnetic beads (100 μL) were washed twice with RIPA buffer (supplemented with protease inhibitor cocktail), then the beads were incubated with an equal volume of lysate from the respective genotype overnight at 4°C. The next day, the lysate was discarded and the beads washed twice with 1 mL of RIPA buffer (supplemented with protease inhibitor cocktail), once with 1 mL of 1 M KCl, once with 1 mL of 0.1 M Na_2_CO_3_, once with 1 mL of 1 M urea in 10 mM Tris-HCl, pH 8.0, and twice more with 1 mL of RIPA buffer (supplemented with protease inhibitor cocktail). For the western blot, the RIPA buffer was removed, protein loading buffer was added (1X Laemmli loading buffer supplemented with 2 mM biotin and 20 mM DTT) to the beads, and the samples were heated at 95°C for 10 min to elute the proteins for further analysis.

## LC-MS

### Protein identification and quantitation by liquid chromatography–mass spectrometry (LC–MS/MS) using label-free data-independent acquisition (DIA)

The relative abundance of proteins was determined by label-free relative quantitative proteomics (MaxLFQ) using data-independent acquisition (MaxDIA) (*77*). Proteins were purified and concentrated by a brief SDS-PAGE run designed to focus all proteins into a single band of about 1 cm x 0.5 cm. Gels were washed 3 times in double-distilled water for 15 min each. Proteins were visualized and fixed by staining with EZ-Run Protein staining solution (ThermoFisher Scientific, USA). The stained protein gel regions were excised and de-stained 3 times in 50% methanol for 15 min each time, followed by washes with 50 mM NH_4_HCO_3_ in 30% acetonitrile until completely de-stained. Finally, each gel sample was dehydrated with 100% acetonitrile. Cysteinylated proteins were reduced with 10 mM dithiothreitol and alkylated with 55 mM iodoacetamide. After two washes with 50 mM NH_4_HCO_3_ in 30% acetonitrile, samples were dried by vacuum centrifugation. Dried gel samples were digested overnight with 500 ng mass spectrometry-grade trypsin (Trypsin Gold, Promega, Madison, WI, USA) in 50 mM NH_4_HCO_3_ digest buffer. After acidification with 10% formic acid to a final 1% formic acid, peptides were extracted with 5% formic acid/50% acetonitrile (v/v) and concentrated to a small droplet using vacuum centrifugation. Desalting of peptides was performed using hand-packed SPE Empore C18 Extraction Disks as described (*78*). Desalted peptides were again concentrated and reconstituted in 10 μL of 0.1% formic acid in water.

Aliquots of the peptides were analyzed by nano-LC-MS/MS using an Easy nLC 1000 equipped with a self-packed 75 μm × 20 cm reverse phase column (ReproSil-Pur C18-AQ, 3 μm, Dr. Maisch GmbH, Germany) coupled online to a Q Exactive HF Orbitrap mass spectrometer via a Nanospray Flex source (all instruments were from Thermo Fisher Scientific, Waltham, MA, USA). Analytical column temperature was maintained at 50°C by a column oven (Sonation GmbH, Germany). Peptides were eluted with a 3%–40% acetonitrile gradient over 110 min at a flow rate of 250 nL/min. The mass spectrometer was operated in data-independent acquisition mode. MS survey scans were acquired in profile mode at a resolution of 120,000 (at m/z 200) over a scan range of 300–1650 m/z. Following the survey scans, 30 groups of precursors were selected for fragmentation with sliding isolation windows to include peptide m/z values ranging from 364 to 1370 Th. The default maximum charge state was set to 4, and resolution was set to 30,000. In MS/MS, the fixed first mass was set to 200 Da. The normalized collision energy (NCE) /stepped NCE was 25.5, 27, and 30. The maximum injection time for the survey scan was 60 ms, and for MS/MS, it was set to auto. The ion target value for both scan modes was set to 3 × 106. Data were analyzed by MaxQuant software (version 2.5.1.0), termed the MaxDIA analysis type. To identify peptides, we used *Drosophila* in silico-generated spectral libraries that were created from the UniProt *Drosophila* protein sequence database (downloaded on 11/14/2025; 13,821 entries plus additional fasta of isoforms) from the Max Planck Institute of Biochemistry data-sharing drive (https://datashare.biochem.mpg.de/s/qe1IqcKbz2j2Ruf?path=/DiscoveryLibraries). To determine the relative abundance of a protein across a set of samples, we used its label-free quantitation (MaxLFQ) intensity calculated by MaxDIA (*79*), which normalizes the protein intensity based on peptide intensities across samples.

### Statistical analysis of mass spectrometry data

We compared gene-level log2-transformed intensity values between the two experimental groups (*Elav>2H1-turboID+α-syn* vs *Elav>Dv^sdAb^-turboID+α-syn*, n=3 replicates per group) using a Welch’s t-test. For each gene, log2 intensities from replicates (Intensity_*Elav>2H1-turboID+α-syn* and Intensity *Elav>Dv^sdAb^-turboID+α-syn*) were used directly without additional transformation. For missing values, we used the Perseus statistical platform (version 2.0.7.0) imputation to replace missing values from a normal distribution at a width setting of 0.3 and a downshift of 1. For each protein, we calculated the mean intensity per group and the log2 fold change (difference between group means on the log2 scale) as an effect size. Raw p-values were adjusted for multiple testing using an artificial variance setting s=0.1 to control the relative importance of t-test, p value and difference between means 4 (*80*). At s=0, only the p values matter whereas at s=0.1, the difference of means plays a role. Results shown in Table S2 include p-values, log2 intensities, and log2 fold changes for downstream analysis.

Significantly changing proteins between *Elav>2H1-turboID+α-syn* and *Elav>Dv^sdAb^-turboID+α-syn* triplicates, with fold differences of <0.67 and >1.5, at p values <0.05, were subjected to STRING Gene Ontology functional enrichment analysis (*81*).

### Seahorse analysis

For Seahorse analysis, we adapted the protocol available from previously published reports (*82, 83*), with minor modifications as described here. The heater on an XFe24 Seahorse Analyzer was turned off 24 h before conducting *Drosophila* experiments to allow it to cool to near room temperature. A fresh sensor cartridge’s Port A was loaded with 56 µL of AHL media containing 25 µM of oligomycin (final concentration 2.5 µM; manufacturer’s protocol), and Port B with 69 µL of adult hemolymph liquid [AHL media-energy source: 10 mM glucose + 10 mM Na-pyruvate + 10 mM Na-lactate + 1 mM glutamine; buffer: 4 mM HEPES; ions: 108 mM NaCl, 5 mM KCL, 8.2 mM MgCl_2_, 1 mM NaH_2_PO_4,_ pH 7.3] containing 5 µM of rotenone (final concentration 0.5 µM; manufacturer’s protocol). The analyzer was calibrated with this cartridge for about 1 h before loading the *Drosophila* brains. After dissection, the brains were placed into an XFe24 Spheroid plate containing 500 μL of AHL media per well, then carefully positioned at the bottom center of each well using forceps. The analyzer was set to cycle through mix/wait/measure for 1 min/0 s/3 min. Baseline OCR was measured over 7 cycles, followed by 6 additional cycles for oligomycin and 7 cycles for rotenone injection. Sudden OCR fluctuations during mixing, caused by tissue movement away from the sensor probe, were observed, and those wells were excluded from the analysis.

### Mammalian expression and purification of sdAbs

For culture studies, epitope mapping and BLI assays, the sdAb clones were expressed in a mammalian expression vector pVRC8400-sdAb. These constructs were made by inserting their individual gene sequences between the 5′ Eco RI and 3′ Afe I sites with a signal peptide, mouse interleukin-2 leader sequence (MYRMQLLSCIALSLALVT) at the N terminus, and myc-his tags at the C terminus for detection and purification. Briefly, Freestyle 293F cells (Invitrogen, catalog no. R790–07) were transfected with the mixture of DNA plasmid and polycation polyethylenimine (PEI; 25 kDa linear PEI, Polysciences Inc., catalog no. 23966) and then incubated at 125 rpm, 37°C, with 5% CO_2_ in Freestyle 293 expression medium. Supernatants were harvested 5 days after transfection, filtered through a 0.45-μm filter, and then purified using Ni–nitrilotriacetic acid (NTA) columns (GE Healthcare). Following elution, each sdAb was dialyzed into PBS, and its concentration was determined by a bicinchoninic acid (BCA) assay.

### Epitope mapping of the sdAbs against *α*-synuclein peptide libraries using the dot blot assay

The sdAbs 2H1, 2H7, and 1D12 were tested on a dot blot to examine their binding to an α-syn peptide library (synthesized by Genscript Inc., Paramus, NJ), aimed to identify their binding epitopes. The library included all 140 amino acids of α-syn, with peptides 1-13 each containing 15 amino acids and peptide 14 containing 10 amino acids, with five amino acid overlaps between peptides. For epitope mapping, peptides were dissolved in a small amount of dimethyl sulfoxide and diluted to 1 mg/mL in PBS. Each peptide (5 μg) was dot-blotted onto a nitrocellulose membrane and air-dried for 30 min. The membrane was blocked for 1 h with 5% milk in 0.1% Tween-20 in TBS-T, then incubated overnight at 4°C with sdAbs 2H1, 2H7, and 1D12 (containing his tag, at 0.01 mg/mL) in Superblock (Thermo Fisher Scientific). After washing several times with TBS-T, the blots were incubated for 1 h at room temperature with rabbit anti-6×His tag antibody (1:2000, Abcam, EPR20547) in Superblock. Following additional TBS-T washes, signals were detected with an IRDye 800CW secondary antibody (1:10,000, LI-COR Biosciences). The bands were quantified using LI-COR ImageStudio Lite 5.2.

### Biolayer Interferometry (BLI) binding affinity assay

To determine the binding affinity of sdAbs for peptides 7, 8, and 9, we measured K_D_ values using BLI (ForteBio Octet RED96). BLI experiments were performed at room temperature in a 96-well black flat-bottom plate (Greiner Bio-One, 655209) with shaking at 1000 rpm. sdAb samples were prepared in 200 μl of assay buffer (1× PBS, 0.05% Tween-20, pH 7.4).

### Solid phase

An SA biosensor was used to assess biophysical interactions between the sdAb 2H1 and biotinylated α-syn (rec α-syn) prepared with EZ-Link NHS-PEG4-Biotin. Then, 1 mg of rec α-syn in 0.5 mL of PBS (pH 7.4) was combined with a 10 mM solution of EZ-Link™ NHS-PEG4-Biotin in DMSO, and the mixture was gently rotated at room temperature for 30 min. Excess biotin was then removed through dialysis. Before starting BLI, the SA biosensor was hydrated in assay buffer for 10 min, and a 120 s baseline was recorded. Next, an optimized concentration of biotin-labeled α-syn preparation (typically 400 nM for rec α-syn) was loaded for 120 s. A second 120 s baseline was recorded, followed by association for 300 s at various 2H1 concentrations and dissociation for 400 s in assay buffer. A reference biosensor was loaded with biotinylated α-syn and tested in an assay buffer blank during the association and dissociation phases. Biosensor tips were regenerated by cycling three times for 10 s each between 10 mM glycine (pH 2) and assay buffer.

### Solution phase

The Ni-NTA biosensor was used to assess the biophysical interaction between the his-tagged sdAbs and the peptides. The sensors were pre-immobilized with nickel-charged tris-nitrilotriacetic groups to capture his-tagged molecules. The sensors were hydrated in the assay buffer for at least 10 min before running the assay protocol. Following this, a baseline step was performed in the assay buffer for 120 s. Next, the Ni-NTA tips were loaded with his-tagged sdAb at 5 μg/ml for 120 s, allowing specific binding between his tag and nickel ions. A subsequent baseline measurement was then performed in the assay buffer for another 120 s. Afterward, ligand-loaded biosensor tips were immersed in solutions of α-syn at various concentrations in PBS (pH 7.4) containing 0.05% Tween-20 (PBS-T) for 300 s. Dissociation was then monitored in the assay buffer for 400 s. A reference biosensor was prepared with his-tagged sdAb and measured with an assay buffer blank during association and dissociation. To regenerate the biosensor tips, they were cycled three times for 5 s between 10 mM glycine (pH 2) and assay buffer, followed by a 1-min recharging step with 10 mM NiCl_2_.

Data analysis was conducted using Data Analysis 11.0 software, applying reference subtraction based on a 1:1 binding model as described previously in detail (*15*).

### Microscopy and documentation

Tissues were imaged using a Zeiss LSM700 confocal microscope at consistent settings. Identical optical/digital parameters, along with an equal number of optical sections, were used to construct the final projection images in ImageJ. Fluorescence intensity was calculated in ImageJ and plotted in GraphPad Prism version 10.6.1. Finally, Adobe Photoshop CS5 was used to assemble images of different genotypes for comparison.

### Statistics

For each figure, error bars in the graphs are presented as mean + or ± SEM, and the data were analyzed using GraphPad Prism version 10.6.1. Climbing assay readings were analyzed using a two-way ANOVA followed by Tukey’s multiple-comparisons test. Data from cell culture, western blotting, immunostaining, and mitochondrial assays were analyzed by a one-way ANOVA followed by either Tukey’s or Dunnett’s multiple-comparisons test. The survival assay was analyzed by the Logrank (Mantel-Cox) test. RT-PCR data were analyzed by an unpaired t-test.

## Acknowledgements

We thank Juan Botas at the Department of Molecular and Human Genetics, Baylor College of Medicine, Houston, Texas, the Bloomington Drosophila Stock Center, and the Developmental Studies Hybridoma Bank (DSHB) for fly stocks and reagents. We are grateful to the NDRI (Philadelphia, PA), from which we purchased a human patient brain with extensive cortical α-syn inclusions, which is a hallmark of LBD. This work was partially supported by National Institutes of Health grants R56AG083436, R01AG032611, R01NS077239 to EMS, RF1/R01NS120488 to EMS and HDR, and S10RR027990 to TAN.

## Author contributions

Pragati conducted most of the fly studies, related analysis, and binding affinity/epitope mapping experiments with different α-syn peptides presented in this study. EC performed cell culture experiments. LCMS data was run and analyzed by HE-B and TN with help from Pragati, HDR, and EMS. The binding affinity experiments with recombinant α-syn in the solid and solution phases were performed by Y.J. The anti-α-syn sdAb transgenic lines were generated by H-WH with help from ISM. The sdAbs for in vitro studies were expressed and purified by RP. and X-PK. All experiments and initial drafts were designed and written by Pragati, HDR, and EMS. All authors had the opportunity to edit the article. HDR and EMS supervised the project.

## Data and materials availability

All data needed to evaluate the conclusions in the paper are present in the paper and/or the Supplementary Materials. Proteomics LC-MS/MS data (raw mass spectrometry data, peak lists and results) that support the findings of this study are available at MassIVE (UCSD) (https://massive.ucsd.edu/ProteoSAFe/static/massive.jsp) under the deposition number MSV000102218 and are deposited to the ProteomeXchange Consortium via the MassIVE partner repository and can be retrieved with the dataset identifier PXD079930.

## Supplementary Figures

Figures S1 to S14

## Other Supplementary Material

Tables S1 to S2

---

**Figure S1.**
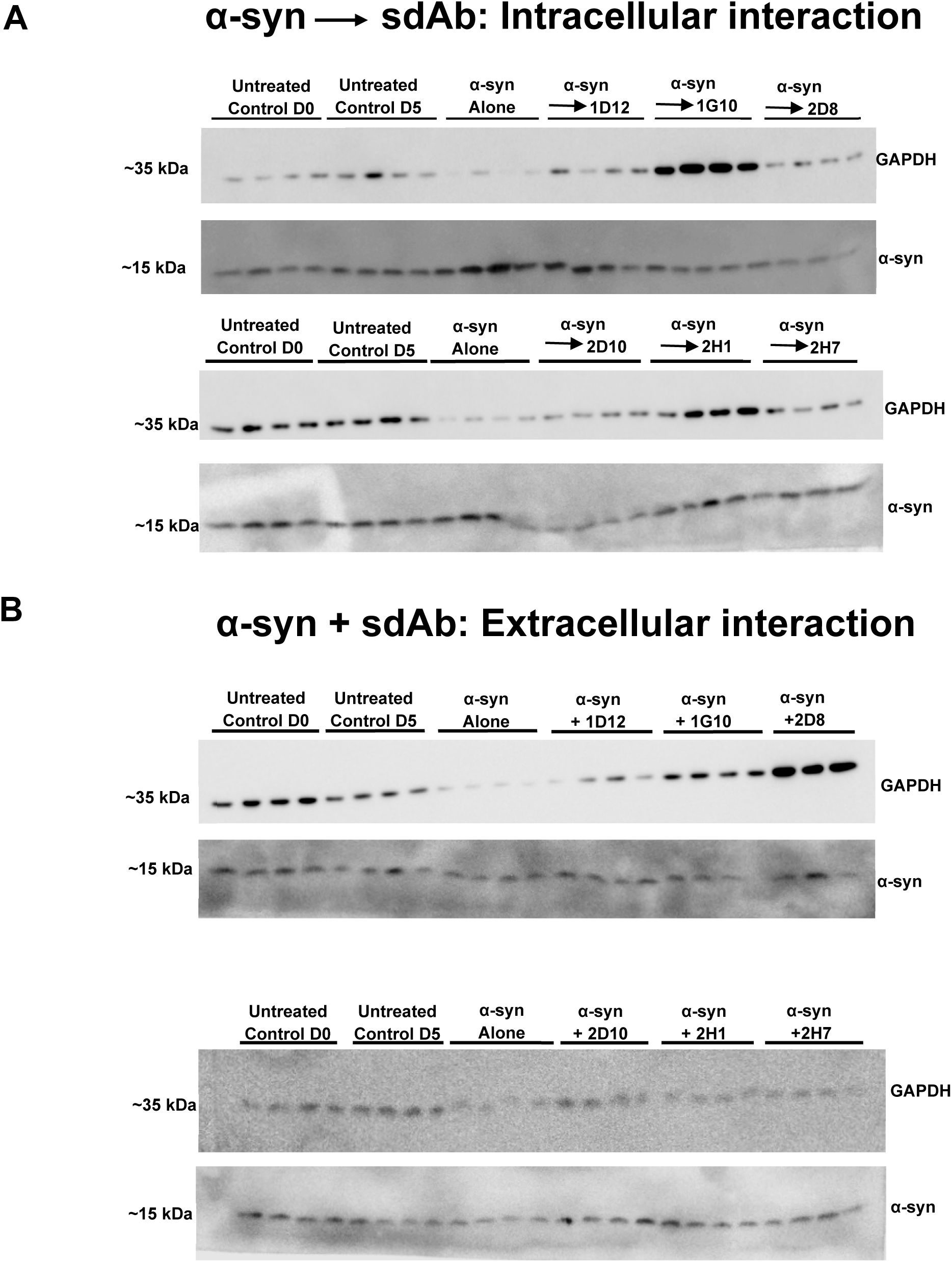
Representative western blots used for quantification in. **Fig. 1**. (A-B) Immunoblots of GAPDH and α-syn levels in treated M83 cell lysate of intracellular (A) and extracellular (B) paradigms. In culture, all sdAbs except one (1D12) prevented α-syn aggregation while some prevented its toxicity.

**Figure S2:**
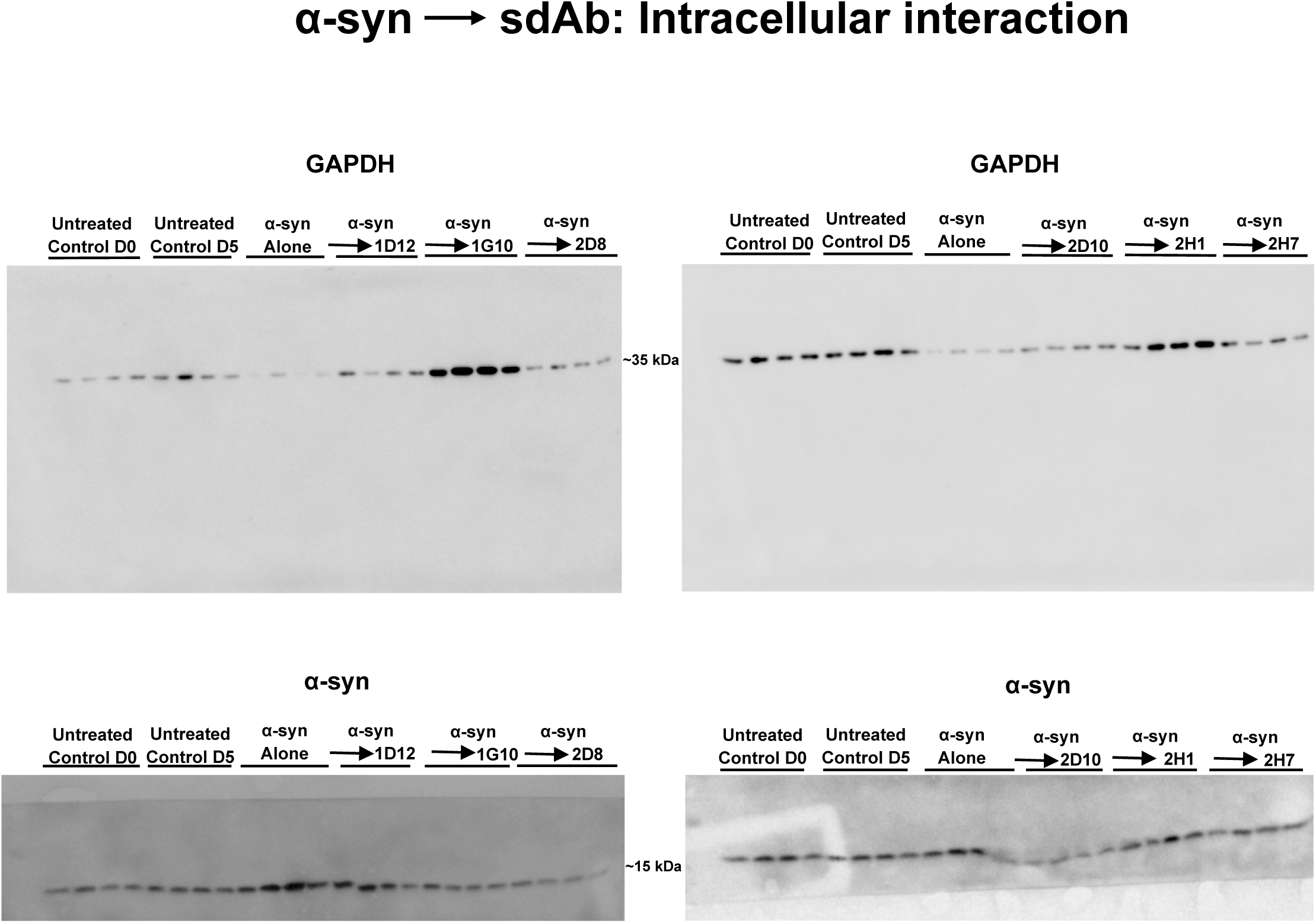
Complete western blots of representative blots shown in Fig. S1A. Some of the blots were cut to allow them to be reacted with different antibodies recognizing proteins at different molecular weights. Molecular weight marker was added to the first well of each gel (Precision Plus Dual Color Standards, BioRad). These bands are visible in the membrane upon transfer. The blots were developed using a chemiluminescent agent that reacts to the secondary antibody but does not always visualize the protein ladder.

**Figure S3:**
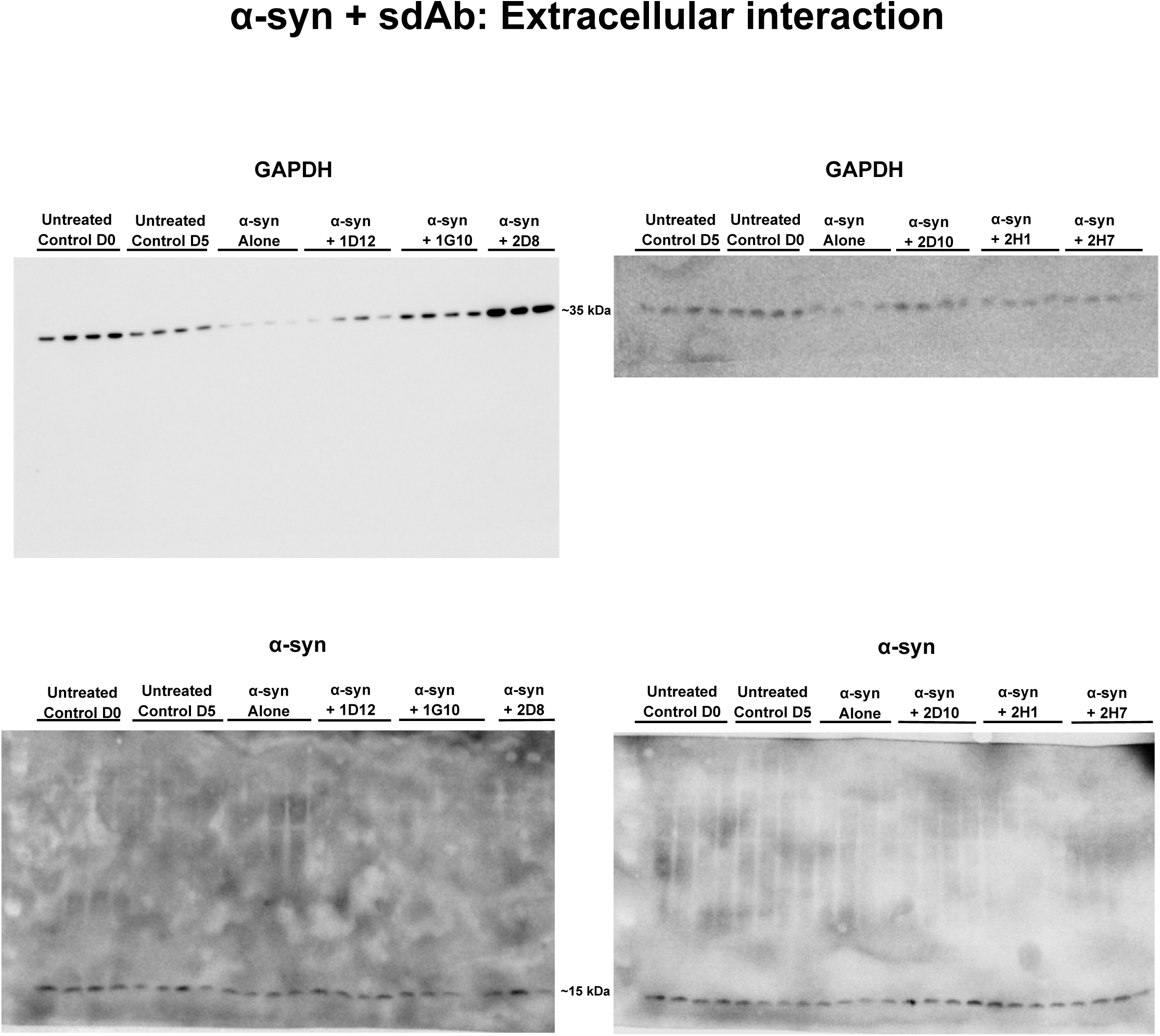
Complete western blots of representative blots shown in Fig. S1B. Some of the blots were cut to allow them to be reacted with different antibodies recognizing proteins at different molecular weights. Molecular weight marker was added to the first well of each gel (Precision Plus Dual Color Standards, BioRad). These bands are visible in the membrane upon transfer. The blots were developed using a chemiluminescent agent that reacts to the secondary antibody but does not always visualize the protein ladder.

**Figure S4:**
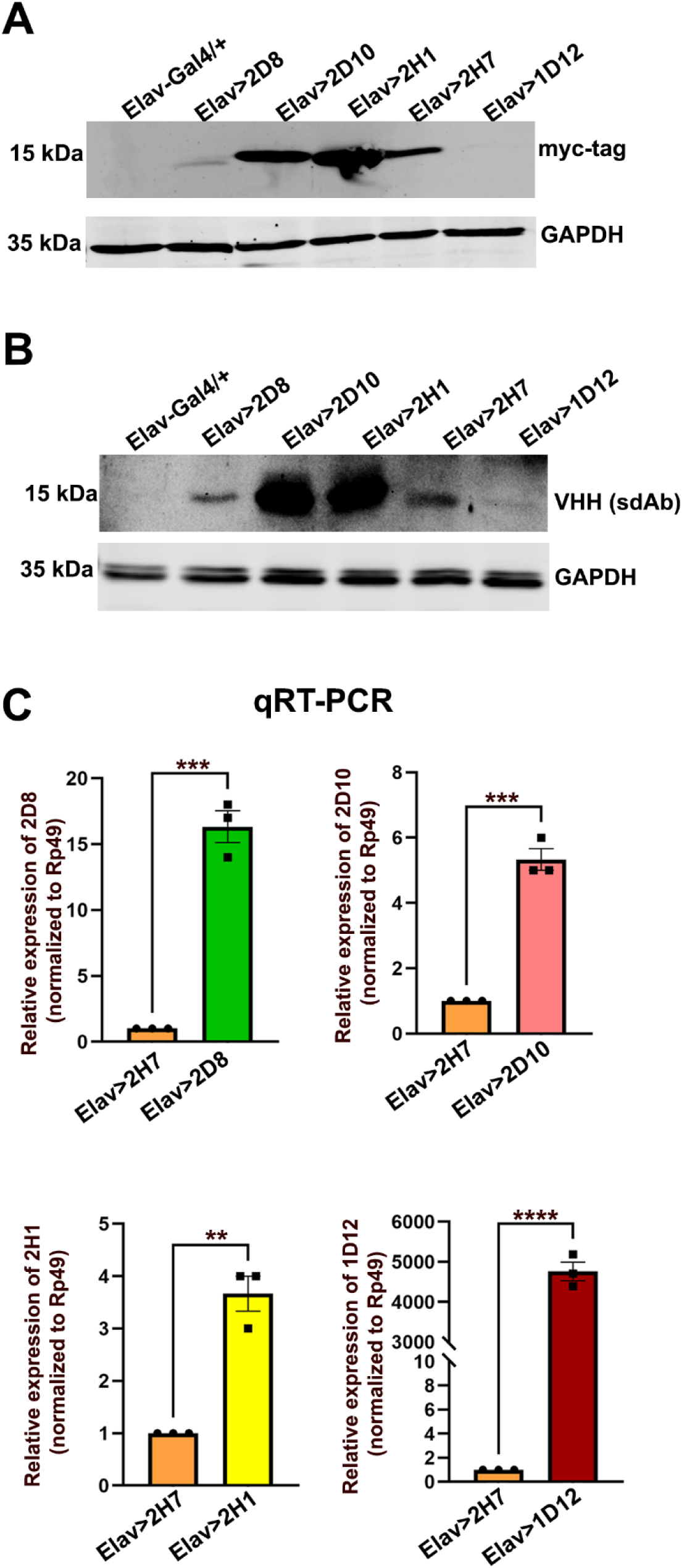
Pan neuronal expression of anti-*α*-syn sdAbs in females. (A-B) Immunoblot of protein levels of different anti-α-syn sdAbs from 5-day-old female flies detected with antibodies to myc-tag, VHH, and a GAPDH loading control. (A) Myc antibody detected robust sdAb expression in Elav>2H1, Elav>2D10, and Elav>2H7, low sdAb expression in Elav>2D8, and did not detect a signal in Elav>1D12. (B) VHH antibody detected strong expression in Elav>2D10 and Elav>2H1, followed by Elav>2H7, Elav>2D8, and Elav>1D12. (C) Quantitative RT-PCR of 5-day-old adult female fly heads driven pan-neuronally. All the sdAbs (2D8, 2D10, 2H1, 2H7, and 1D12) were clearly transcribed, with 1D12 having the highest transcript abundance. Relative expression was normalized to the housekeeping gene RP49. Bar graphs are presented as mean ± SEM. Unpaired t-test, two-tailed. **p ≤ 0.01, *** p ≤ 0.001, and ****p ≤ 0.0001.

**Figure S5:**
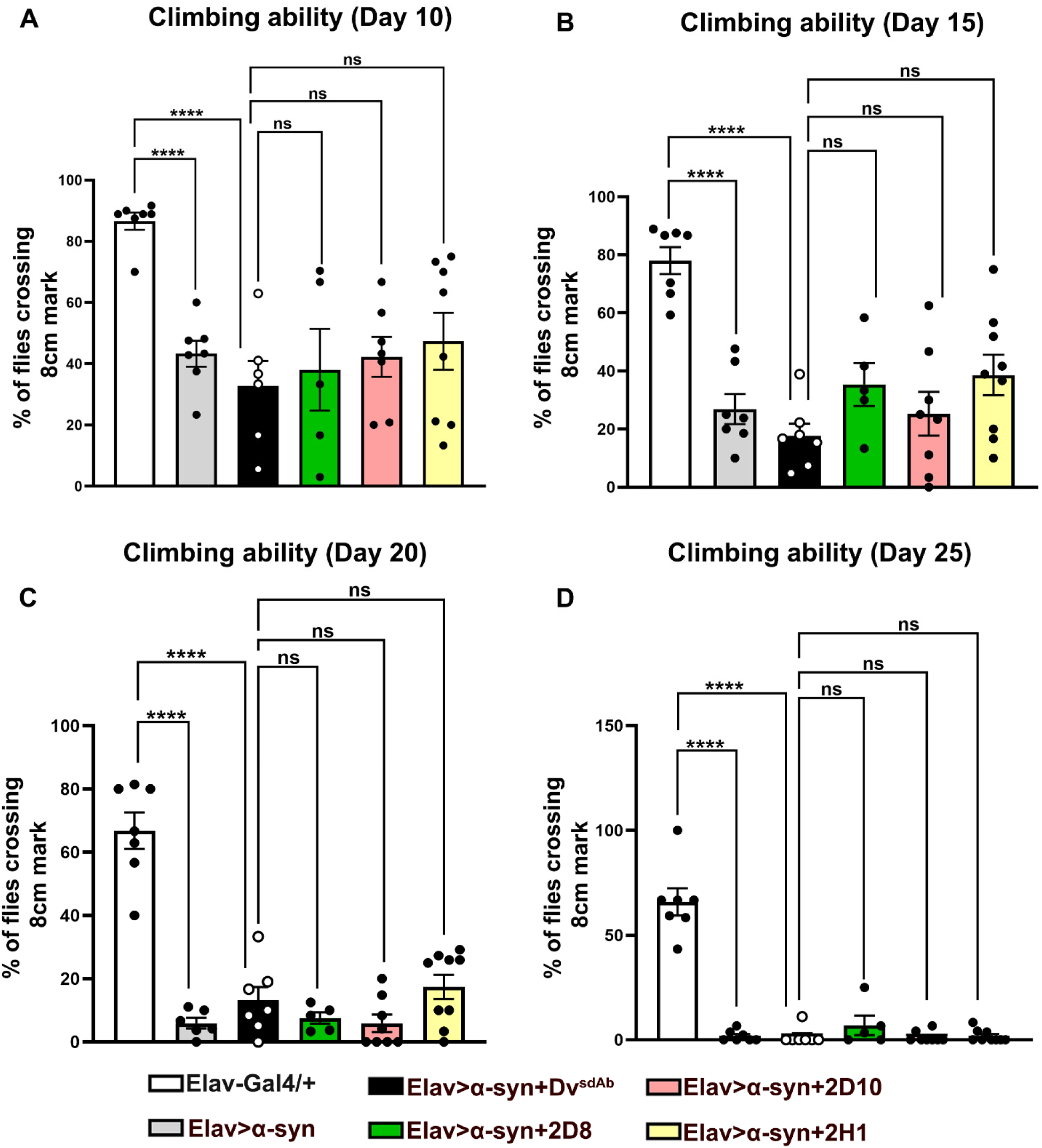
Pan-neuronal expression of anti-*α*-syn sdAbs does not rescue climbing defects in *α*-syn-expressing male flies. Quantitative assessment of the relative climbing ability of different age-matched adult male flies. Results from day 10, day 15, day 20, and day 25 are shown. Different genotypes are indicated by distinct colors as shown in the figure legend. Flies that do not express α-syn (white bars) climbed faster than those that do express α-syn. The grey and black bars represent flies expressing α-syn and α-syn with a control sdAb (Dv^sdAb^). These two groups exhibited impaired motor function at all time points, and their condition worsened with age. None of the three anti-α-syn sdAbs improved climbing ability, compared to the control Dv^sdAb^ group. Bar graphs are presented as mean ± SEM, and each data point represents an average of 5-10 flies in a vial. Genotypes and the number of flies analyzed per group: *Elav-Gal4/+* (N = 57), *Elav>α-syn* (N = 97), *Elav>α-syn+Dv^sdAb^*(N = 63), *Elav>α-syn+2D8* (N = 42), *Elav>α-syn+2D10* (N = 69), and *Elav>α-syn+2H1* (N = 78). Two-way ANOVA, Tukey multiple-comparison test. ****p ≤ 0.0001 and ns = non-significant.

**Figure S6:**
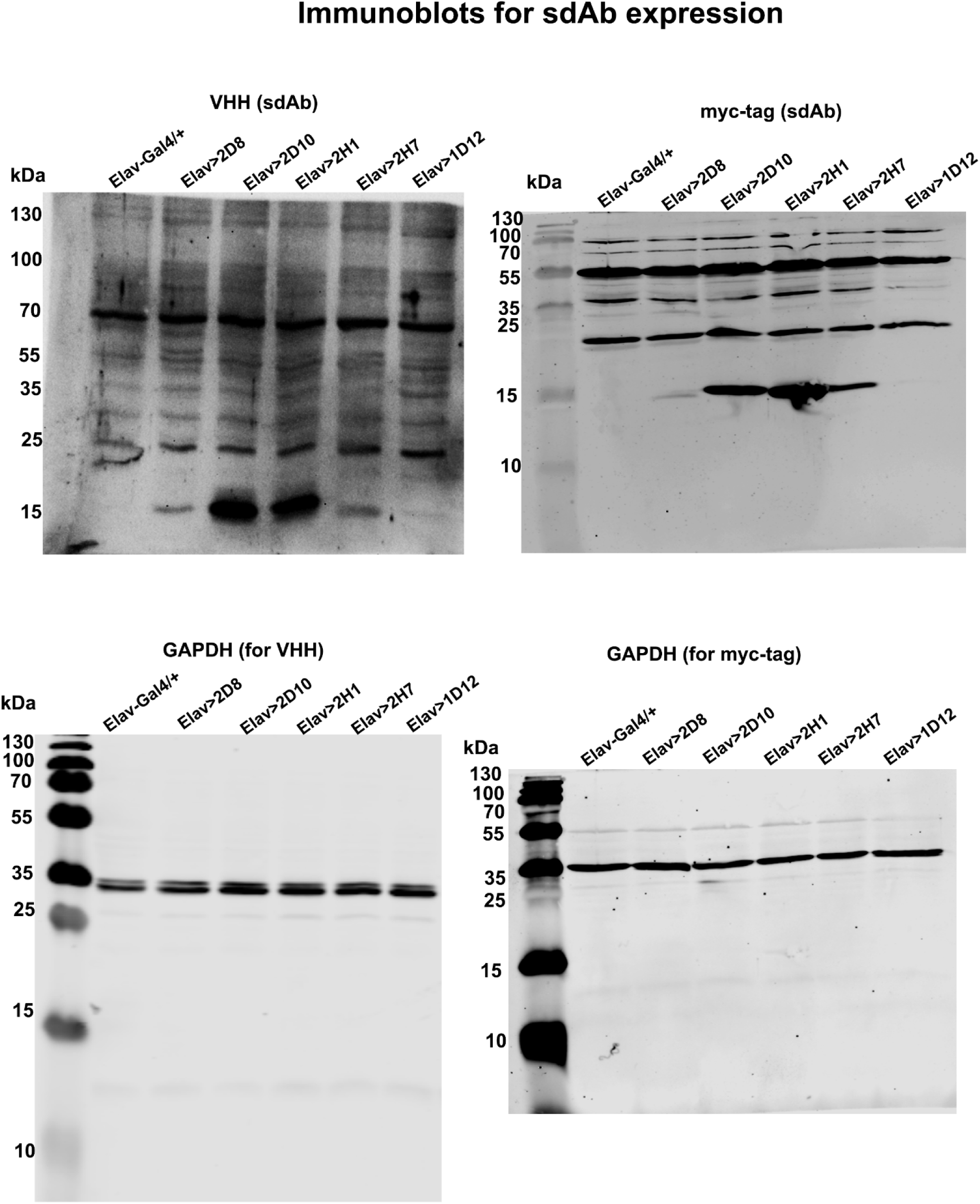
Uncropped western blots of representative blots shown in Fig. S4.

**Figure S7:**
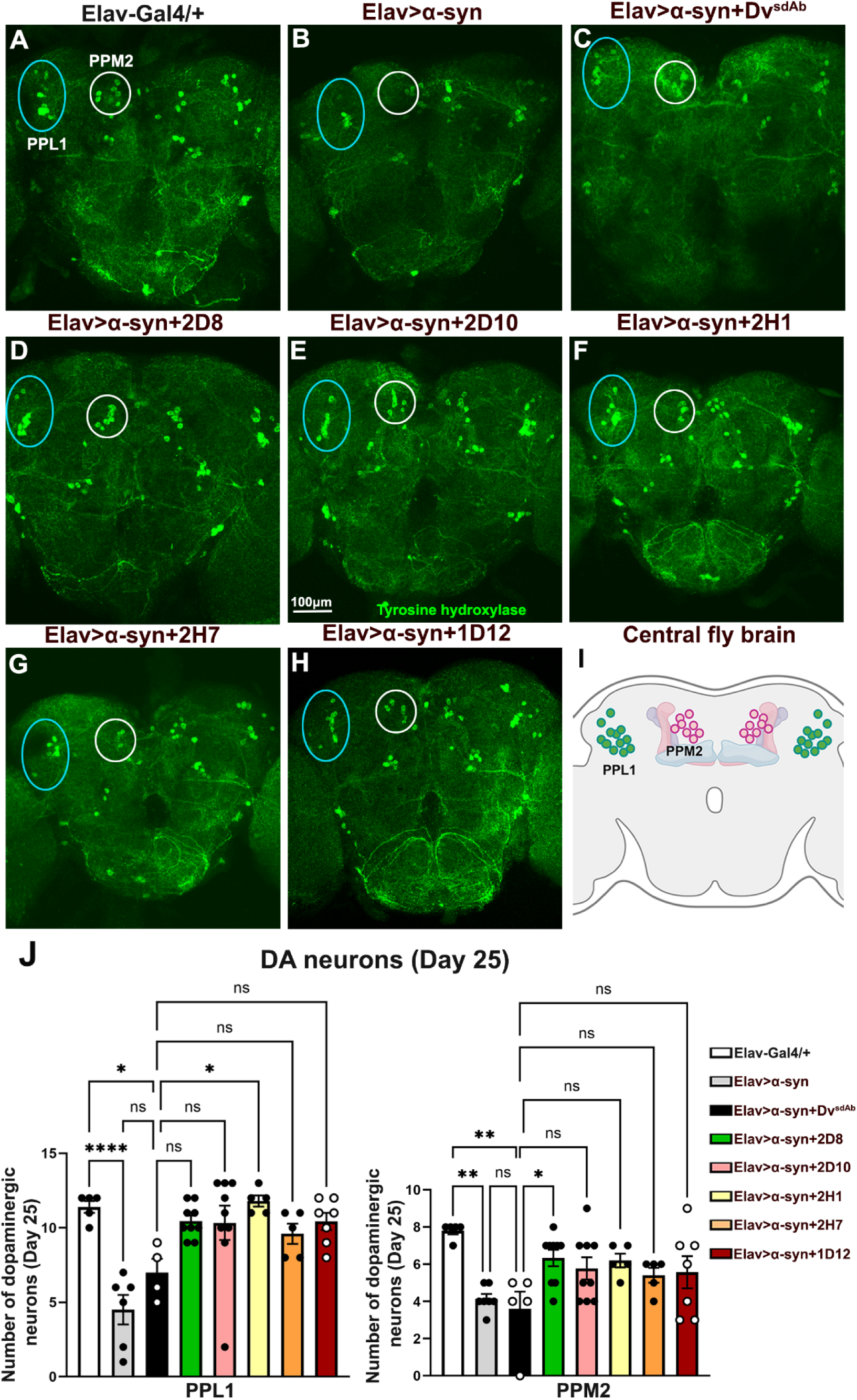
Pan-neuronal expression of anti-*α*-syn sdAbs prevents the loss of DA neurons in *α*-syn-expressing flies. (A-H) Fluorescence images of 25-day-old female adult brains stained with anti-tyrosine hydroxylase antibody. The marked circles outline PPL1 (blue) and PPM2 (white) neuronal clusters. (I) The schematic shows PPL1 and PPM2 neuronal clusters in an adult fly brain. (J) The average number of DA neurons in PPL1 and PPM2 neuronal clusters in different age-matched female adult brains per hemisphere (N = 5-9 per genotype). *Elav>α-syn* and *Elav>α-syn+Dv^sdAb^*(control sdAb) flies had loss of DA neurons, and sdAb 2H1 prevented loss of PPL1, whereas sdAb 2D8 prevented the loss of PPM2 DA neurons in *Elav>α-syn* flies, whereas all other sdAbs were ineffective at 25 days post eclosion. Bar graphs are presented as mean ± SEM. One-way ANOVA, Tukey post hoc test. *p ≤ 0.05; **p ≤ 0.01, ****p ≤ 0.0001 and ns = non-significant; Scale bar A-H = 100µm.

**Figure S8:**
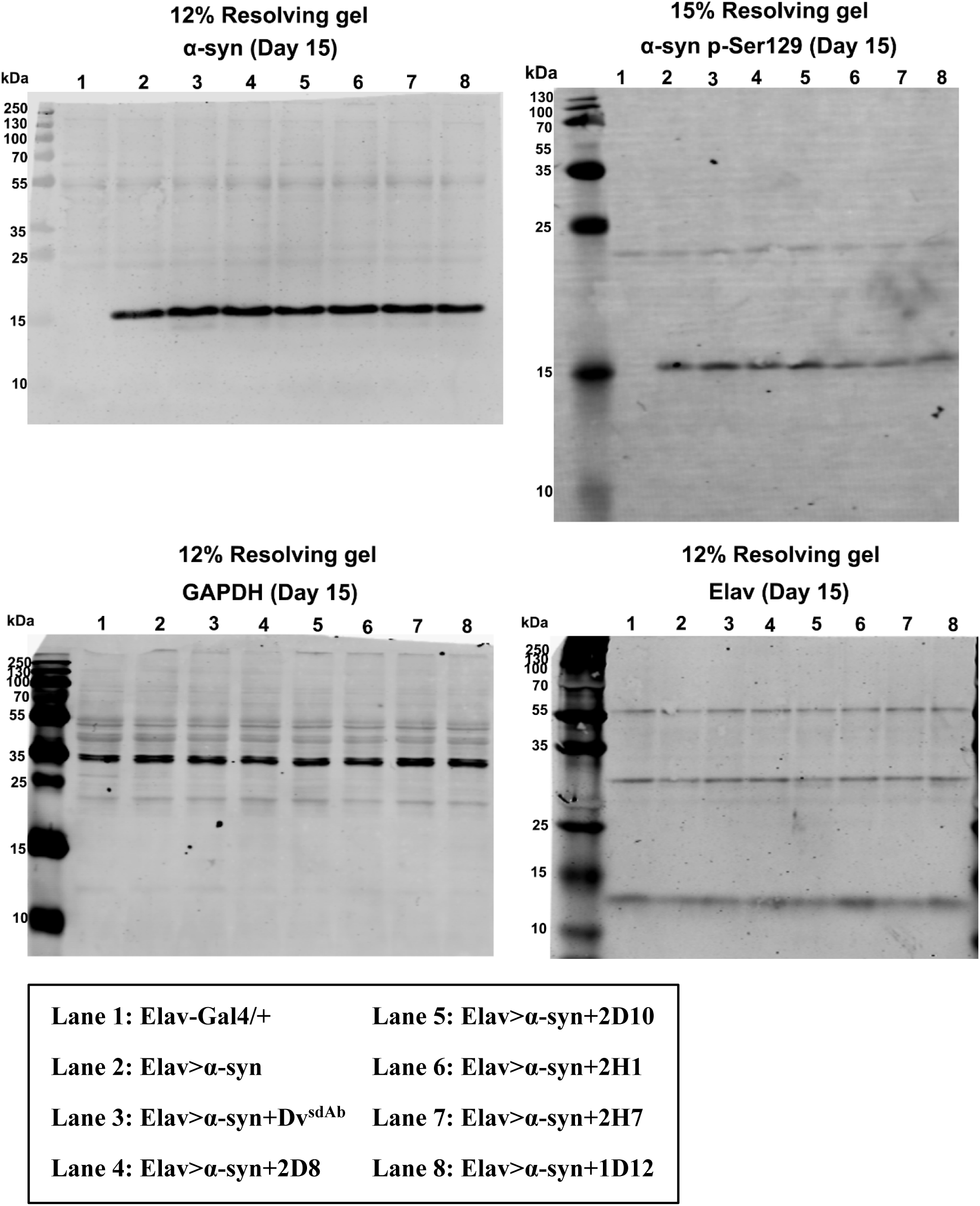
Uncropped western blots of representative blots shown in Fig. 3A.

**Figure S9:**
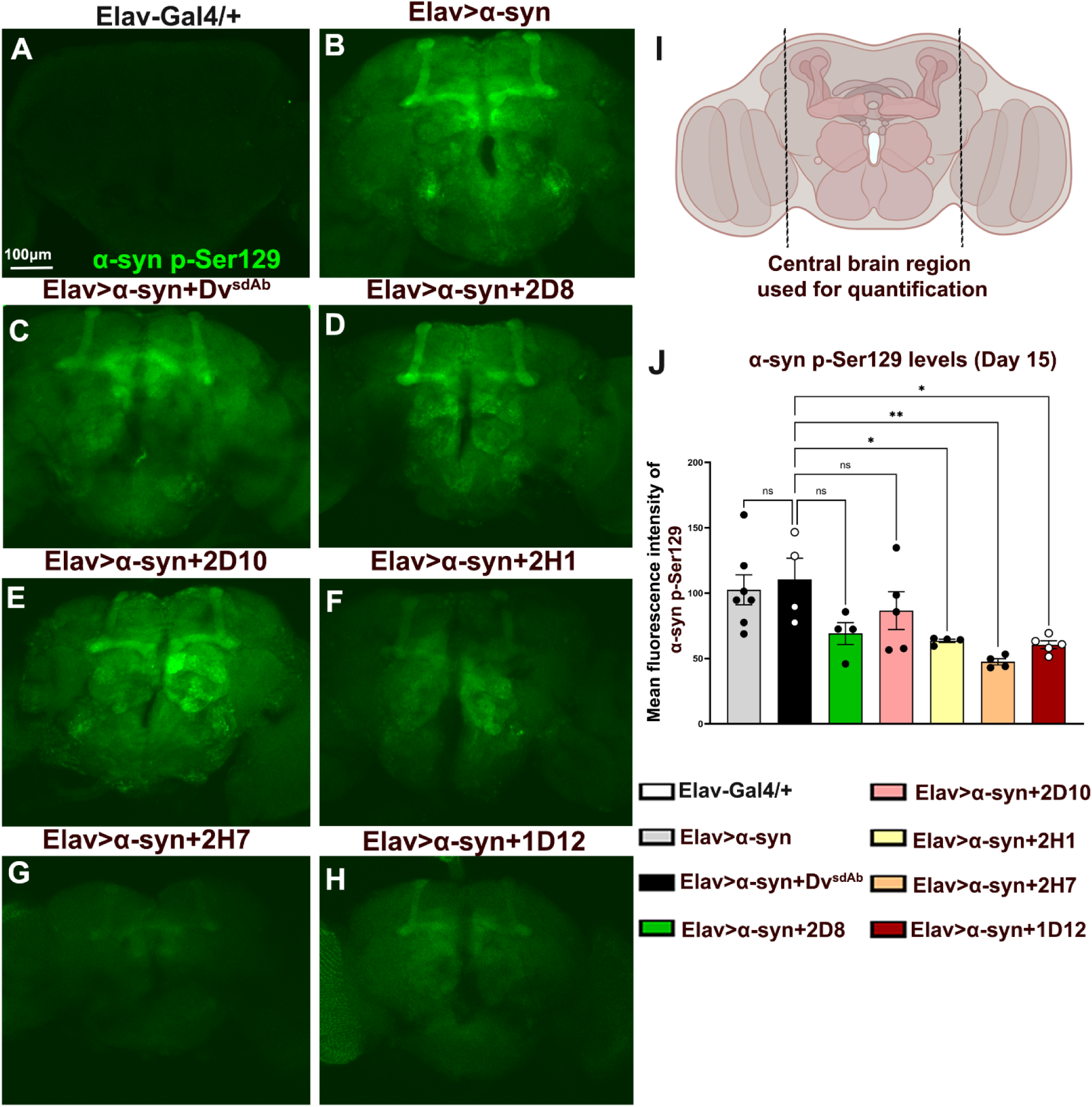
Neuron-specific expression of anti-*α*-syn sdAbs reduces pathological *α*-syn p-Ser129 immunoreactivity in brains of adult female flies expressing *α*-syn in neurons. (A-H) Confocal images of 15-day-old female adult brains stained with α-syn p-Ser129 antibody. *Elav-Gal4/+* control flies were immunonegative for α-syn p-Ser129 (A). (I) The schematic shows the central brain region of an adult fly brain. (J) Quantification of α-syn p-Ser129 immunoreactivity in the central brain region marked in I. sdAbs 2D8 and 2D10 did not reduce α-syn p-Ser129, whereas sdAbs 2H1, 2H7, and 1D12 decreased α-syn p-Ser129 levels (N = 4-7 per genotype). Scatter plot bar graphs are presented as mean ± SEM. One-way ANOVA, Dunnett’s multiple comparisons test. *p ≤ 0.05, **p ≤ 0.01 and ns = non-significant; Scale bar A-H = 100µm.

**Figure S10:**
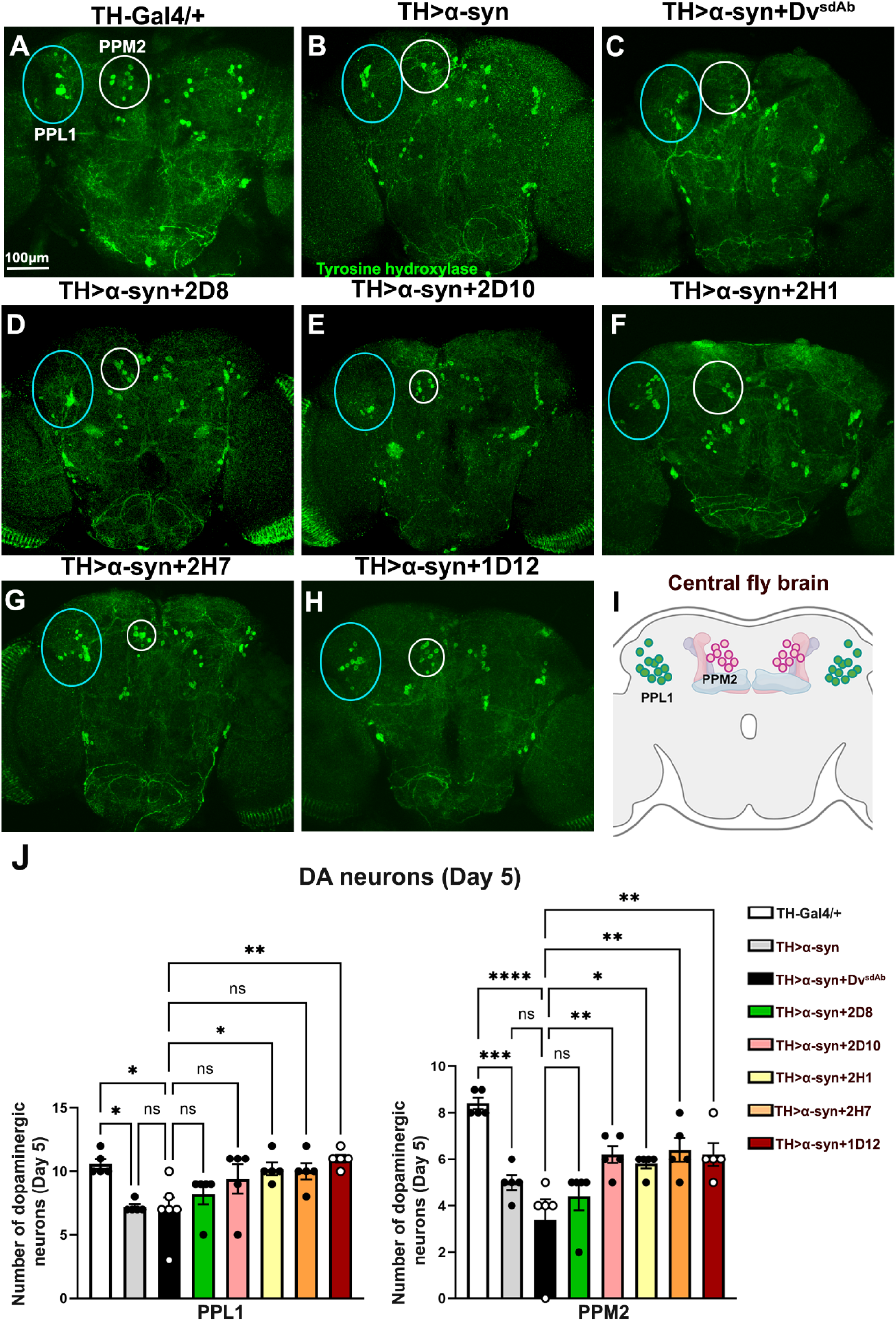
Targeted expression of anti-*α*-syn sdAbs in DA neurons expressing *α*-syn prevents loss of DA neurons. (A-H) Confocal images of 5-day-old female adult brains stained with anti-tyrosine hydroxylase antibody. (I) The schematic shows PPL1 and PPM2 neuronal clusters in an adult fly brain. (J) The average number of DA neurons in PPL1 and PPM2 neuronal clusters in different age-matched adult female flies per hemisphere (N = 5-6 per genotype). *TH>α-syn* and *TH>α-syn+Dv^sdAb^* (control sdAb) flies had loss of DA neurons in different clusters. sdAbs 2H1 and 1D12 prevented loss of PPL1 DA neurons in *TH>α-syn* flies, whereas all sdAbs except 2D8 were effective in preventing PPM2 neuronal loss. Bar graphs are presented as mean ± SEM. One-way ANOVA, Tukey’s multiple-comparison test. *p ≤ 0.05, **p ≤ 0.01, *** p ≤ 0.001, ****p ≤ 0.0001 and ns = non-significant; Scale bar A-H = 100µm.

**Figure S11:**
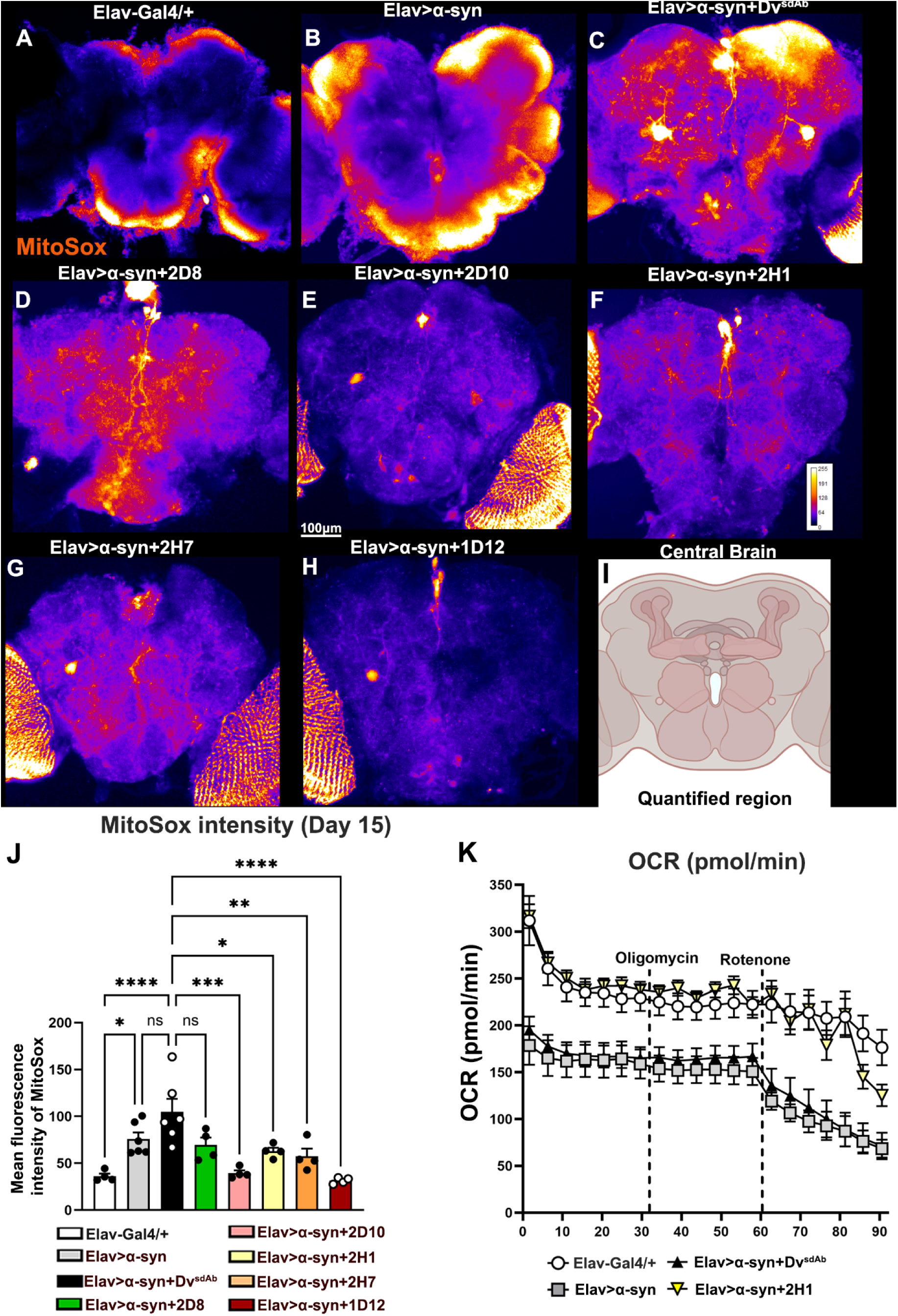
Anti-*α*-syn sdAbs reduce superoxide levels in the mitochondria and improve OCR in synucleinopathy flies. (A-H) Fluorescent images of 15-day-old female adult brains stained with MitoSox Red dye. (I) The schematic shows the central brain region of an adult fly brain. (J) The mean fluorescence intensity of MitoSOX measured in the adult brain shows that sdAb 2D8 was not effective, while sdAbs 2D10, 2H1, 2H7, and 1D12 significantly reduced superoxide levels in α-syn expressing flies (N = 4-6 per genotype). (K) Changes in the OCR levels measured after oligomycin and rotenone administration in adult brains of different genotypes (N=6-8 per genotype). 2H1 improves those levels in α-syn expressing flies to those seen in normal control flies. See Fig. 7 for a condensed version of this data with statistical analysis. Graphs show mean + or ± SEM. One-way ANOVA, Tukey’s multiple comparisons test. *p ≤ 0.05; **p ≤ 0.01, *** p ≤ 0.001, ****p ≤ 0.0001 and ns = non-significant.

**Figure S12:**
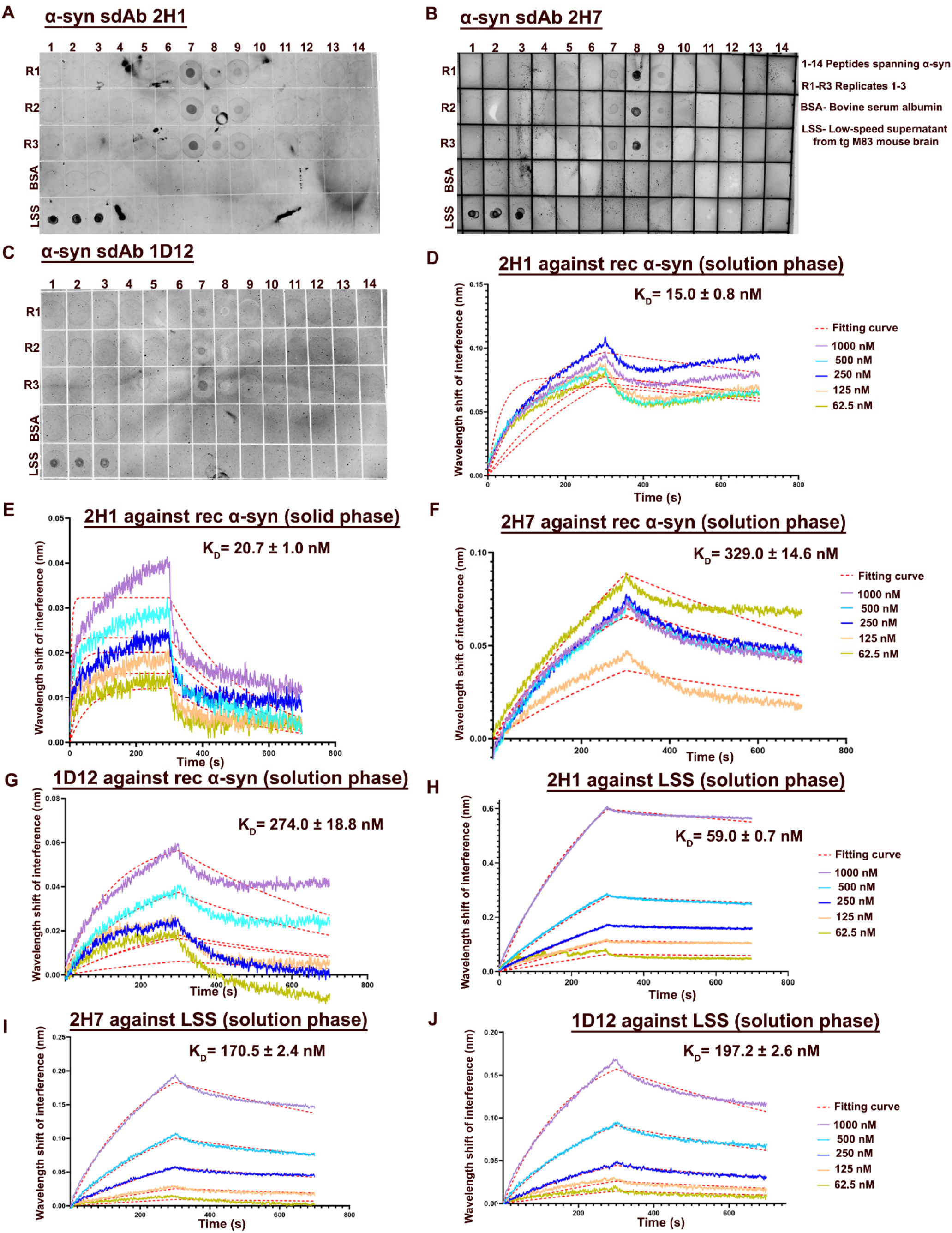
Epitope mapping and binding affinities of anti-*α*-syn sdAbs 2H1, 2H7, and 1D12 by dot blot and BLI binding affinity assay. (A-C) Dot blots of binding of sdAbs 2H1 (A), 2H7 (B), and 1D12 (C) with overlapping 14-amino-acid peptides spanning the entire α-syn protein. Negative control: bovine serum albumin (BSA). Positive control: low-speed supernatant (LSS) fractions. (D-J) Line curves of the wavelength shift of interference (in nm) for different sdAbs 2H1, 2H7, and 1D12 with recombinant α-syn (D-G) and LSS (H-J). For peptide sequences, see Table S1.

**Figure S13:**
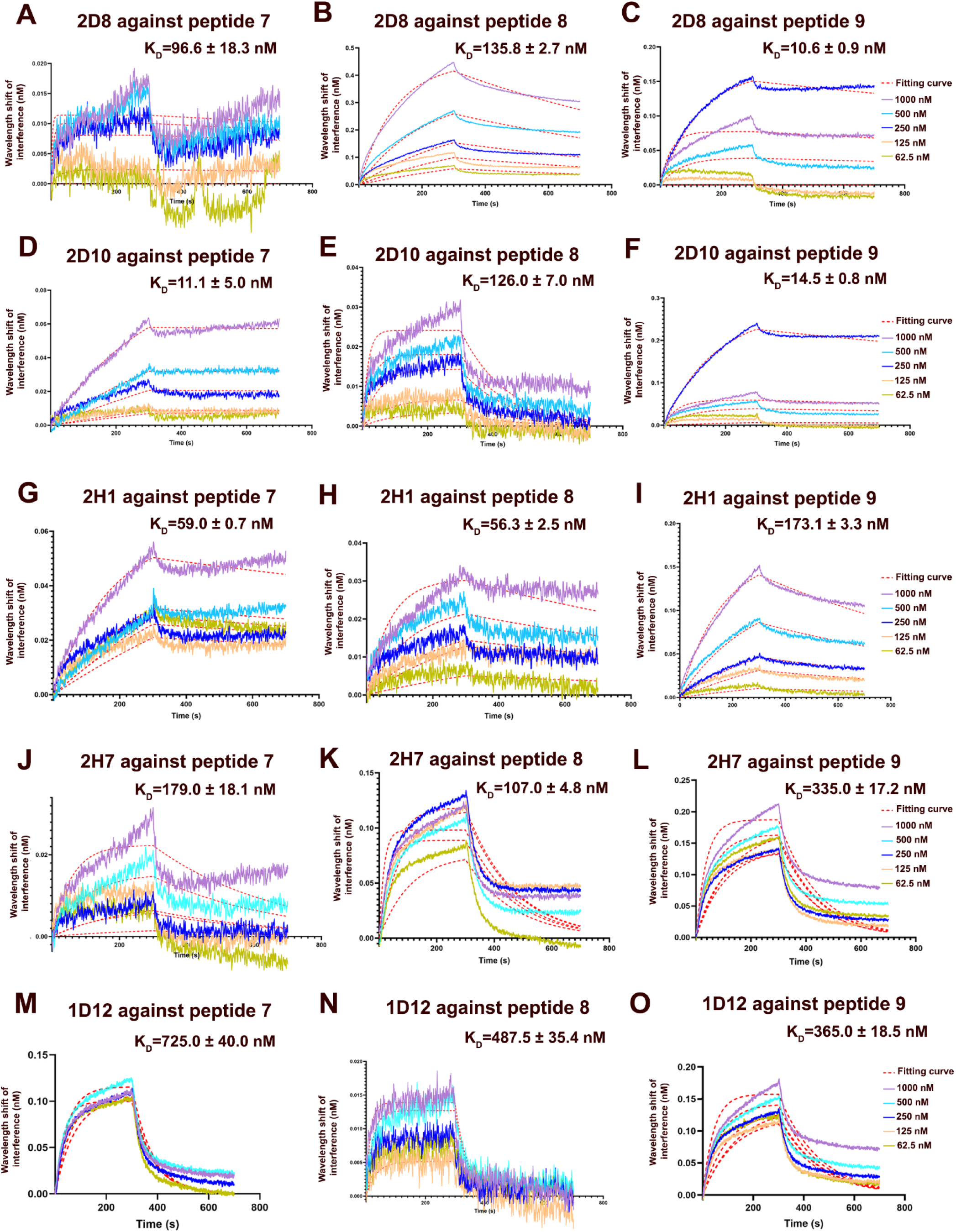
Binding affinities of all anti-*α*-syn sdAbs against peptides 7, 8, and 9 measured in solution phase by biolayer interferometry assays. (A-O) The curves illustrate the wavelength shift of interference (in nm) for different sdAbs with peptides 7, 8, and 9, indicating binding affinity. The curves illustrate the association and disassociation of sdAb with different α-syn peptides at various concentrations. The red line is the fitting curve used to determine the KD value ± standard deviation (SD) from three independent experiments.

**Figure S14:**
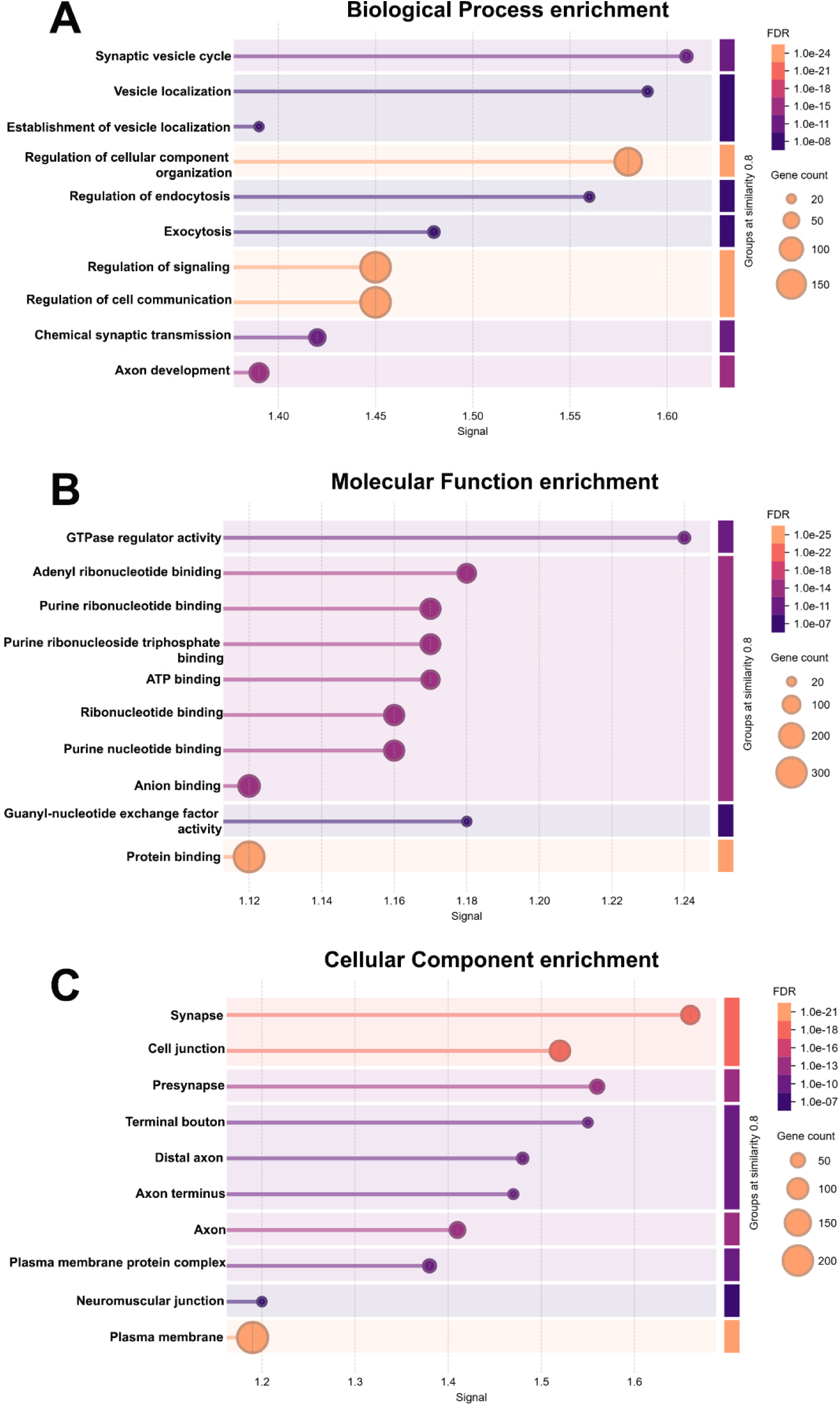
Biological process, molecular function, and cellular components of enriched proteins in *Elav>2H1-turboID+α-syn* flies. Pathway analysis indicates enrichment of (A) synaptic vesicle cycle, (B) GTPase regulator activity, and (C) synapse as prominent biological processes, molecular functions, and cellular components, respectively, in *Elav>2H1-turboID+α-syn* flies.

**Table S1: :**
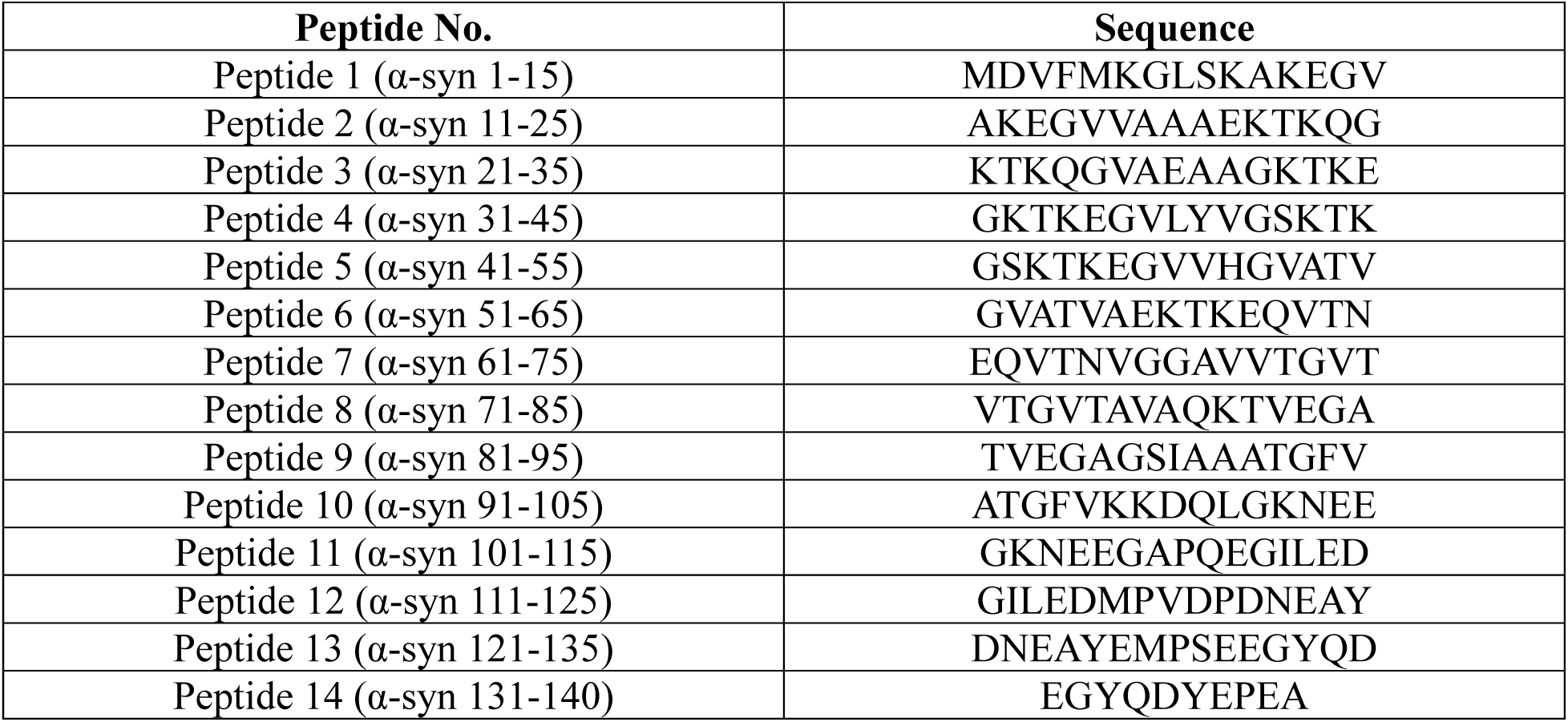
Sequences of *α*-synuclein peptide library used for epitope mapping. To mimic the charge state in the native protein, peptides 2-14 are acetylated on the N-terminus and peptides 1-13 are amidated on their C-terminus.

